# Insights into Ligand-Mediated Activation of an Oligomeric Ring-Shaped Gene-Regulatory Protein from Solution- and Solid-State NMR

**DOI:** 10.1101/2024.05.10.593404

**Authors:** Rodrigo Muzquiz, Cameron Jamshidi, Daniel W. Conroy, Christopher P. Jaroniec, Mark P. Foster

## Abstract

The 91 kDa oligomeric ring-shaped ligand binding protein TRAP (*trp* RNA binding attenuation protein) regulates the expression of a series of genes involved in tryptophan (Trp) biosynthesis in bacilli. When cellular Trp levels rise, the free amino acid binds to sites buried in the interfaces between each of the 11 (or 12, depending on the species) protomers in the ring. Crystal structures of Trp-bound TRAP show the Trp ligands are sequestered from solvent by a pair of loops from adjacent protomers that bury the bound ligand via polar contacts to several threonine residues. Binding of the Trp ligands occurs cooperatively, such that successive binding events occur with higher apparent affinity but the structural basis for this cooperativity is poorly understood. We used solution methyl-TROSY NMR relaxation experiments focused on threonine and isoleucine sidechains, as well as magic angle spinning solid-state NMR ^13^C-^13^C and ^15^N-^13^C chemical shift correlation spectra on uniformly labeled samples recorded at 800 and 1200 MHz, to characterize the structure and dynamics of the protein. Methyl ^13^C relaxation dispersion experiments on ligand-free apo TRAP revealed concerted exchange dynamics on the µs-ms time scale, consistent with transient sampling of conformations that could allow ligand binding. Cross-correlated relaxation experiments revealed widespread disorder on fast timescales. Chemical shifts for methyl-bearing side chains in apo- and Trp-bound TRAP revealed subtle changes in the distribution of sampled sidechain rotameric states. These observations reveal a pathway and mechanism for induced conformational changes to generate homotropic Trp-Trp binding cooperativity.

## Introduction

Ligand-mediated regulation of protein function involves changes in its functional state that accompany binding of one or more ligands [1–3]. When the ligands bind at sites remote from the site of function, this regulation is referred to as “allosteric” [4,5]. Allosteric regulation is widespread in biology, governing processes ranging from trans-membrane signaling, enzyme activity and gene expression[6–9].There is considerable interest in understanding the mechanisms of allosteric regulation, as this could provide means of manipulating essential processes, or engineering new desired regulatory systems [4,5,10,11]. NMR spectroscopy has advantages for studying allostery due to is unique ability to provide information on both structure and dynamics at multiple sites, detecting lowly populated conformational states, and providing site-specific information on conformational entropy [1,12–15]. Of paramount interest is detailing *how* ligand binding at an allosteric site is transmitted through the protein to sites responsible for its activity.

To gain insights into mechanisms of allosteric regulation we studied the ring-shaped undecameric (11-mer) protein TRAP (*trp* RNA binding attenuation protein) from *Geobacillus stearothermophilus* (Figure 1). TRAP is a paradigmatic allosteric protein that regulates expression of the Trp biosynthesis genes at the levels of transcription and translation [16,17]. At high cellular concentrations of tryptophan (Trp) TRAP binds up to eleven Trp ligands and is activated to bind with high affinity to specific sequences in the 5’ leader of the *trp* operon mRNA. This interaction leads to premature transcription termination, and in the full transcript, results in sequestration of the Shine-Dalgarno sequence, inhibiting its translation. At low Trp levels, unbound (apo) TRAP is inactive for RNA binding. Crystal structures of Trp-bound (holo) TRAP reveal that Trp ligands bind between adjacent protomers with the indole ring buried in the hydrophobic pocket formed by β-sheets [18–20]. The Trp ligands bind cooperatively to TRAP, such that successive ligand binding events occur with higher apparent affinity [21,22]. Specificity for Trp arises from a series of polar contacts between each Trp ligand and the BC loop of one protomer and the DE loop of the adjacent protomer [23].

**Figure 1.**
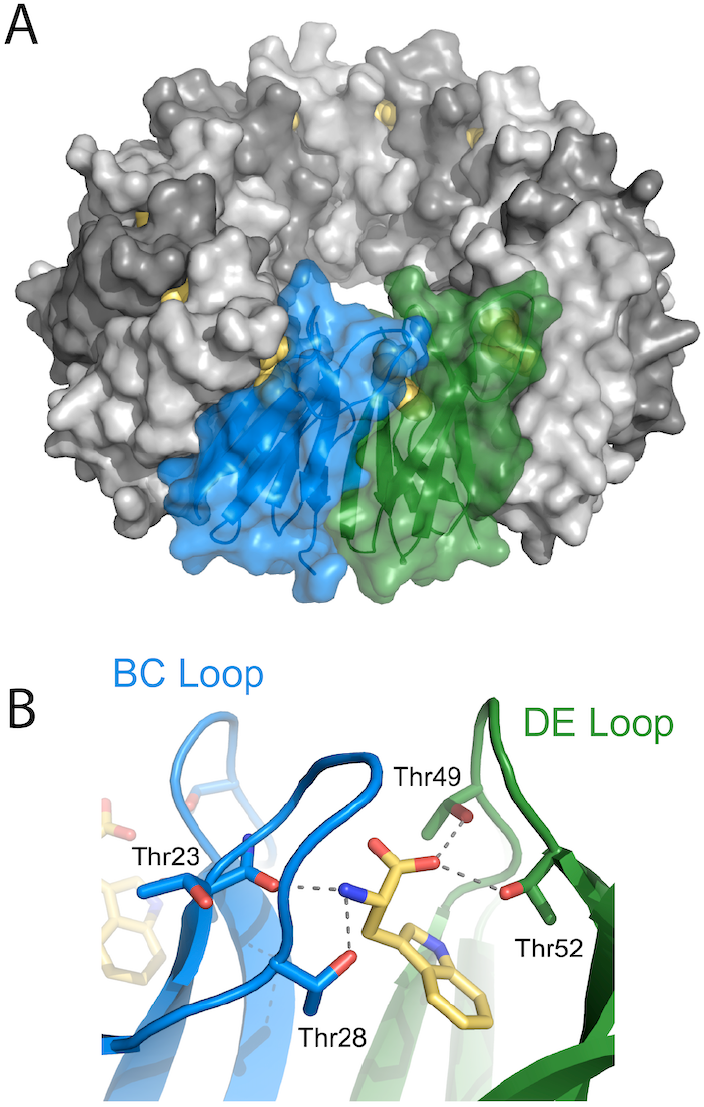
Homo-undecameric TRAP rings bury up to eleven Trp ligands in the interface between adjacent protomers. (A) Undecameric TRAP ring as observed in Trp-bound crystals (PDB:1C9S) shows the ligands are completely excluded from solvent. One protomer is blue, one green, and the others are grey. (B) Four threonine residues from the BC and DE loops contact the carboxyl and amino groups of the buried Trp ligands

Crystal structures show that the resulting binding pocket entirely secludes Trp from solvent. This suggests that in addition to the observed Trp-bound state the protein must sample alternative conformations that allow Trp to bind to and dissociate from its sites in the protein-protein interfaces.

Prior solution NMR studies of the 90.6 kDa TRAP oligomer have shown that in the absence of Trp a high degree of conformational flexibility in the ligand binding loops results in extreme line broadening of backbone amide resonances [24–26]. This flexibility presents obstacles to detailed structural analysis of apo TRAP by both crystallography and solution NMR; that is, the parts of the protein that are of most interest are also the most difficult to observe. The large molecular weight of the protein rings also presents a challenge to solution NMR as their slow molecular tumbling results in fast transverse relaxation, broad signals and decreased intensity in spectra. Despite these limitations, the use of TROSY-based NMR methods along with increased temperature and uniform deuteration enabled backbone resonance assignments of a most of the residues in holo TRAP and 39 residues in apo TRAP; notably, most of the residues in the BC and DE loops could be assigned in holo TRAP but are absent in spectra of apo TRAP [24]. Labeling of the methyl groups of Ile, Leu and Val sidechains enabled more detailed characterization of the effect of ligand binding on the structure and dynamics of the protein, although those residues are not present in the BC and DE loops of the native protein[25]. These spectral differences between apo and holo TRAP reveal that the ligand profoundly rigidifies the ligand binding site, with more modest effects on residues in the structured of the protein ring [24,25]. Nevertheless, the absence in apo TRAP spectra of signals from the Trp-binding BC and DE loops limits the conclusions that can be drawn about the mechanism of Trp access to its sites, or the basis for Trp-Trp cooperativity.

Here we advance our understanding of the structure and dynamics of apo TRAP by application of complementary solution and solid-state NMR approaches. Introduction of threonine methyl labeling has enabled more direct characterization of the structure and dynamics of the BC and DE loops. Moreover, the limitations associated with molecular tumbling and intermediate exchange broadening can in principle be alleviated by the use of magic angle spinning (MAS) solid-state NMR, a robust spectroscopic tool that is not inherently limited by molecular size [27– 29]. Specifically, multidimensional solid-state NMR techniques that rely on polarization transfers based on dipolar and J-couplings can facilitate detailed characterization of structure and conformational dynamics for relatively rigid and highly flexible protein domains, respectively, and have been successfully employed to probe a range of high-molecular weight protein complexes and assemblies including membrane proteins, amyloids, supramolecular machines, viral protein assemblies, and chromatin [30–33]. By combining solution NMR spin-relaxation measurements at 600 and 800 MHz, and MAS solid-state NMR at 800 at 1200 MHz, we quantify the loop dynamics in TRAP that gate access by Trp to its binding sites and illuminate the shifts in the free energy landscape that result in cooperative ligand binding.

## Results

### Methyl-TROSY NMR Spectra Reveal Trp-Induced Structural Perturbations

We prepared samples of A26I-TRAP in which the δ_1_-methyl groups of Ile, and γ_2-_methyl groups of Thr were labeled with ^1^H and ^13^C, while other carbon-bound protons were deuterated and all nitrogen positions enriched with ^15^N; i.e., Thr/Ile-[^13^CH_3_], U-[^2^H,^15^N]-A26I-TRAP. The mutation of Ala26 to Ile was previously engineered to introduce a methyl probe in the BC loop of the protein, and to serve as a metric of Trp binding [25]. Trp and RNA binding experiments, and NMR spectra showed that the A26I mutation did not significantly perturb TRAP structure or function. Although the Thr residues labeled here serve as native monitor probes of the loops, the A26I mutation was retained for consistency with those prior experiments [25].

To monitor Trp-induced conformational changes in TRAP, we acquired 2D ^1^H-^13^C-correlated methyl TROSY-HMQC spectra of A26I-TRAP in its Trp-free apo and Trp-bound holo states (Figure 2A). Axial symmetry of TRAP in these states is evident from the observation of a single set of signals in the ^1^H-^13^C methyl spectra. The benefits of the methyl TROSY effect are evident in the spectra of the 90.6 kDa apo protein as we observe seven Ile (including the extra signal from the A26I mutation) and six Thr methyl signals that have uniform intensities. The ^13^C shifts of four Thr are highly degenerate in apo TRAP (Figure 2A), with four of the six near 21.5 ppm, suggestive of similar time- and ensemble-averaged conformations. In the Trp-bound holo form we observe increased dispersion in the ^13^C dimension as well as variable intensities for the Thr methyl signals, with the sixth signal not observable at 55 °C. Comparing the apo and holo spectra we observed increased chemical shift dispersion for all Thr methyl groups even though only four are proximal to the Trp binding site in the crystal structure of Trp-bound TRAP (Figure 1)[34]. This indicates that structural consequences of ligand binding are not restricted to the immediacy of the Trp binding site, implicating allosteric structural changes.

**Figure 2.**
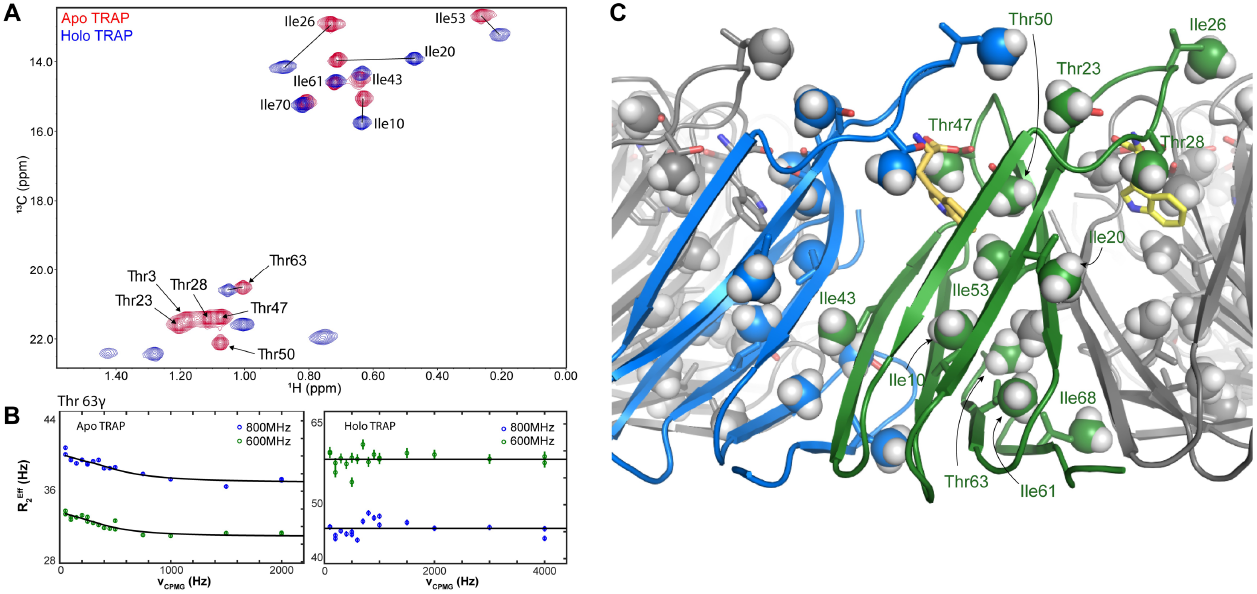
Trp binding alters the structure and dynamics of TRAP rings. (A) ^1^H-^13^C methyl TROSY-HMQC spectra of [U-^2^H/^15^N, Thr/Ile-^1^H/^13^C-methyl]-A26I-TRAP recorded at 800 MHz, 55 °C in the absence (apo, red) and presence (holo, blue) of Trp. Most Thr and Ile methyl resonances are strongly perturbed by Trp binding. (B) Multiple quantum relaxation dispersion (MQ RD) profiles for the Thr63 methyl group recorded at 800 (blue) and 600 (green) MHz at 20 °C in the absence (left) and presence (right) of Trp. Large dispersions observed in spectra of apo TRAP are not observed in spectra of holo TRAP. Lines are fits of the frequency dependence of the effective transverse relaxation rate with a site-specific two-state exchange model for apo TRAP and no exchange for holo TRAP. 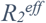 two a site-specific two-state exchange model for apo TRAP and no exchange for holo TRAP.

### Threonine Methyl Resonance Assignments

Thr side chain methyl resonances of TRAP were assigned from a combination of through-space NOE correlations in solution at 55 °C, and through-bond correlations in solid-state NMR spectra. Prior studies had assigned the methyl resonances of Ile, Val and Leu (ILV) residues using a series of solution triple-resonance and “out-and-back” methyl correlation experiments[25]. While those resonances provided useful probes of the structure and dynamics of the apo and holo states of the protein, the absence of native ILV residues in the BC and DE loops left their dynamics poorly understood. In this study Thr methyl signals in apo-TRAP were assigned using 3D CCH HMQC-NOESY-HMQC and ^13^C-edited NOESY-HMQC (HCH) spectra[35]. Five of the six Thr methyls could be assigned based on strong NOEs to proximal Ile methyl groups referencing the crystal structure of Trp-bound TRAP (Table S 1). For example, the Thr50 γ-methyl is within NOE-distance to both Ile53 and Ile26 delta-methyl and shows a cross peak in both the HCH and CCH NOESY spectra (Figure 3, Table S 1). Thr3, present in the disordered N-terminus of the protein was assigned by process of elimination[24].

**Figure 3.**
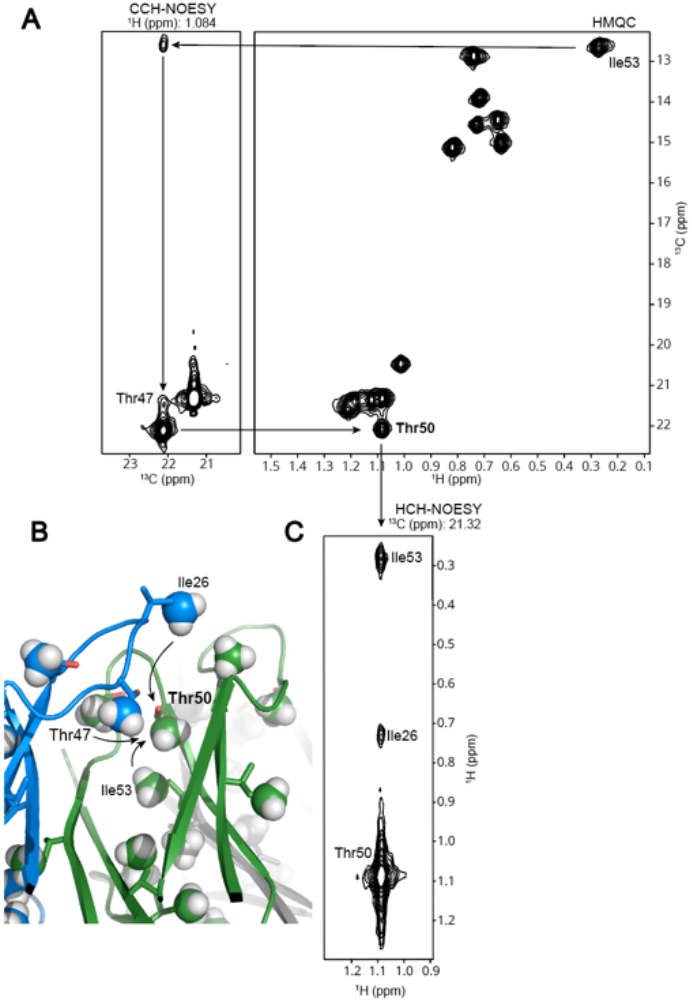
Assignment of Thr γ2 methylresonances of apo TRAP from ^13^C- and ^1^H-separated NOESY spectra, at 55° C. (A) Right, methyl TROSY spectrum; left, CCH-NOESY plane corresponding to the Thr50 γ^2^ methyl ^1^H resonance. (B) Model from the Trp-TRAP crystal structure, illustrating NOEs observed to Thr50. (C) Plane from ^13^C-separated NOESY-HMQC spectrum corresponding to the Thr50 γ^2 13^C methyl resonance.

### CPMG NMR Relaxation Dispersion Reveals Complex Exchange Dynamics

Availability of methyl resonance assignments enabled site-specific interpretation of methyl NMR relaxation dispersion (RD) experiments performed on apo TRAP. RD experiments were performed at 20 °C instead of 55 °C with the expectation that this might result in slow exchange for some of the methyl resonances; methyl spectra at 20°C were of comparable quality (Figure S 1). These experiments revealed dynamics on the µs-ms time scale in the Trp-gating BC and DE loops in apo A26I-TRAP. Multiple-quantum (MQ) ^1^H-^13^C methyl-CPMG (Carr-Purcell-Meiboom-Gill) experiments produce relaxation dispersion (RD) curves in which the effective transverse relaxation rate *R*_2_^eff^ is measured as a function of the frequency at which refocusing CPMG pulses are applied, ν_CPMG_. The resulting values are determined by the intrinsic relaxation rate *R*_2_^0^ and a term *R*_ex_ that quantifies additional relaxation arising from dynamic exchange between states with different chemical shifts that occurs on the time scale of the refocusing CPMG pulse train (µs-ms), *R*_2_^eff^(*ν*_CPMG_) = *R*_2_^0^ + *R*_ex_(*ν*_CPMG_). Fitting the resulting RD curves to a two-state exchange model,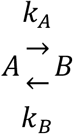, produces values for the rate of exchange between the two states *k*_*ex*_ = *k*_*A*_*+k*_*B*_, their relative populations,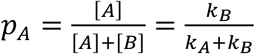, and the chemical shift differences between the exchanging states |Δ*ω*_*C*_ |, |Δ*ω*_*H*_ |. In the fast exchange limit, where *k*_*ex*_ *≫* Δ*ω*, the best-fit |*ω*_*C*_|, |*ω*_*H*_ |, and *p*A values becomes unreliable as many combinations of these parameters can yield equally good fits, and only the *k*_ex_ values can be interpreted[36]. In the case of a global two-site exchange, wherein all signals are experiencing the same exchange process, the RD curves may be “globally fit”, with shared parameters *p*A and *k*_ex_[37].

Measurable relaxation dispersion was observed for all six Thr methyl probes in apo TRAP (Figure 2, Figure 4). All six Thr methyl probes were found to be in the fast exchange regime (*k*_ex_ >>Δ*ω*) and therefore preclude separate quantification of the *p*A and Δ*ω*[38,39]. The RD curves for all six threonine and three of the Ile δ_1_ methyl groups (Ile26, Ile53, Ile61) could be globally fit to a single two-state exchange process with a *k*_ex_ value of 1725 ± 263 s^-1^. Globally fitting the four remaining Ile residues resulted in large residuals so they were fit individually, resulting in unique *k*_ex_ values (1130, 2298, 2088, and 2272 s^-1^).

**Figure 4.**
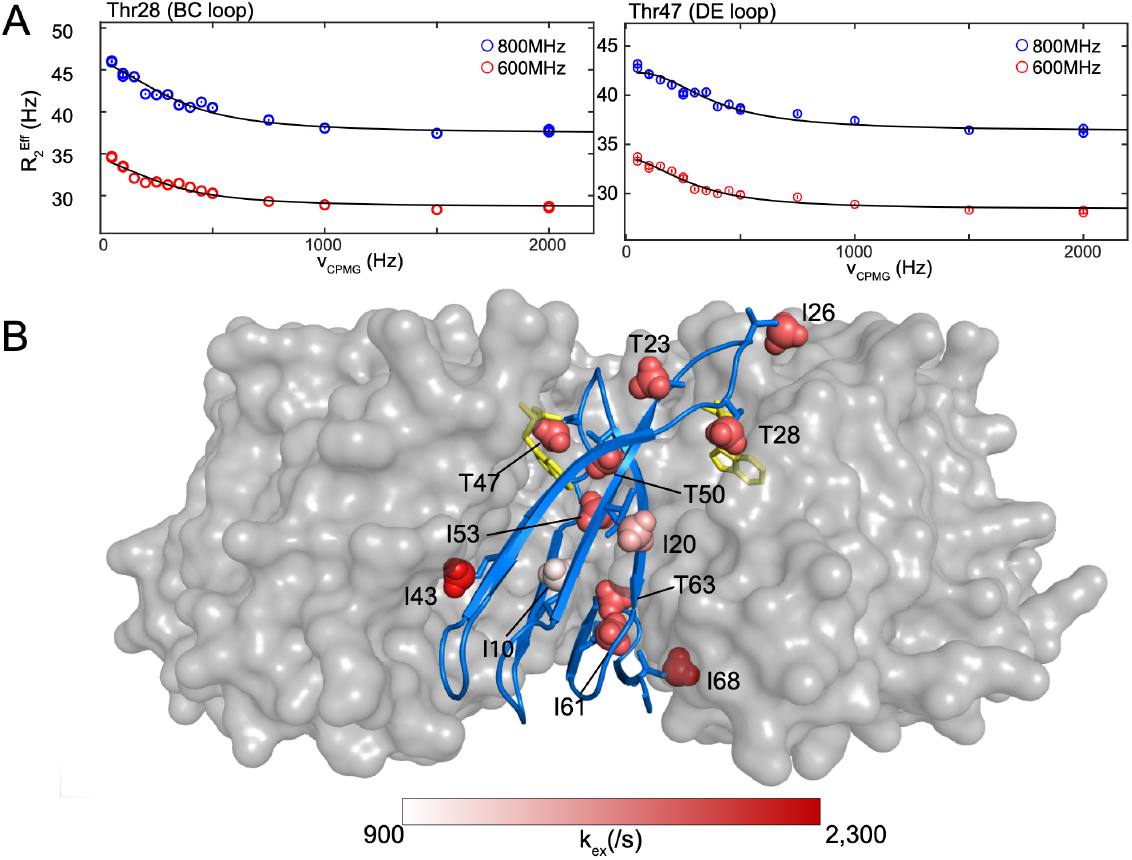
^13^C Methyl relaxation dispersion profiles of apo-A26I-TRAP reveal that all Thr and three of the Ile can be globally fit to a single two-state exchange process, while the other Ile are disparate. (A) MQ RD curves for methyl groups on the BC loop (Thr28, left) and DE loop (Thr47, right) can by fit with the same two-state exchange rate k_ex_ of 1725 ± 263 s^-1^. (B) Methyl k_ex_ values from fitting of MQ RD curves mapped to a single protomer of a model of A26I-TRAP using a linear color ramp from 900 to 2,300 s^-1^ (the max value).

### Methyl Order Parameters

To complement the measurement of µs-ms dynamics in apo-TRAP, we performed methyl dipole-dipole cross-correlated relaxation measurements to quantify the amplitude of fast motion experienced by the methyl bond axis [40,41]. These experiments quantify motion via the generalized order parameter *S*^2^, where a high value (*S*^2^ = 1.0) corresponds to a rigid bond vector on time scales faster than overall tumbling (for TRAP, *τ*_c_ = 32 ns) and a low value (*S*^2^ = 0.0) corresponds to unconstrained motion about the bond vector [42,43]. Intensity ratios for double-quantum and single-quantum coherences were generated from the ^1^H-^1^H dipolar cross-correlated relaxation experiments[42] at 55 °C for all Thr and Ile methyl groups in apo A26I-TRAP and fit with a relaxation rate (*η*) from which we extracted the generalized order parameter (*S*^2^; see Methods) (Figure S 2, Figure S 3). We found that apo-TRAP exhibits large amplitude of motions on the ps-ns timescale as indicated by an average order parameter of ∼0.5 (Figure 5; **Error! Reference source not found**.). Methyl groups from Thr28, Thr47 and Thr50 in the BC and DE loops have low order parameters with *S*^2^ of 0.33, 0.27, and 0.53, compared to an *S*^2^ of 0.99 for Thr63 which is located on β strand F at the core of the protein. This indicates that the Thr residues in the loops are overall more flexible on the ps-ns time scale than those in the rigid core.

**Figure 5.**
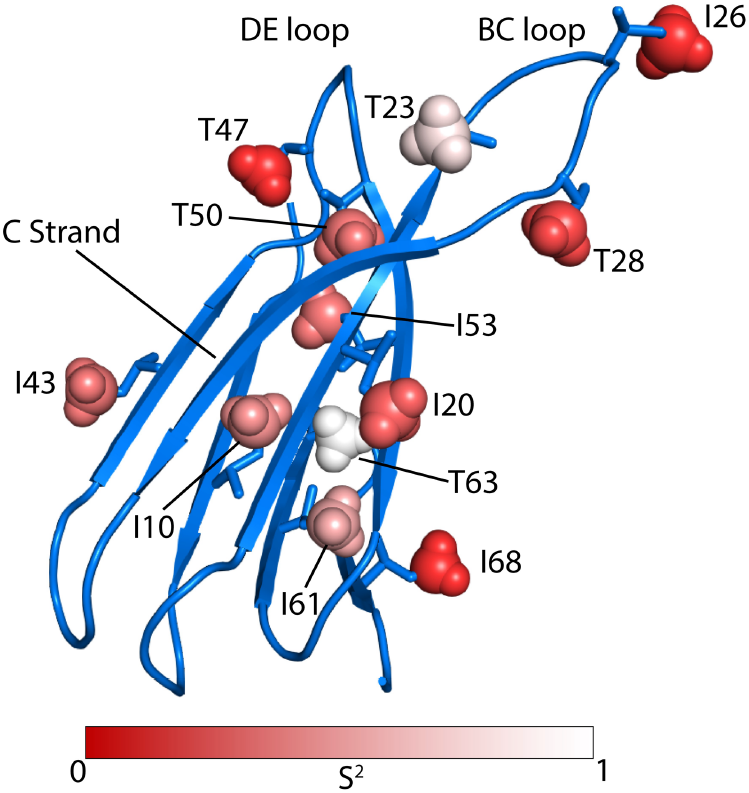
Methyl axis order parameters S^2^ reveal a high degree of conformational flexibility in both the solvent exposed loops and hydrophobic core of A26I-TRAP. Fitted Thr/Ile methyl S^2^ order parameters (Table 1) for mapped to a single protomer of the crystal structure of Trp-bound TRAP. Thr residues in the binding loops exhibit very low S^2^ <0.4. Thr63 exhibits a particularly high order parameter as is anticipated since this residue is in the rigid core of the protein. (Thr3 was not resolved in the crystal structure and is not shown.)

Unexpectedly, Thr23 in the BC loop exhibits a relatively high order parameter of 0.83 despite in a loop predicted to be flexible based the absence of backbone resonances for that region[24]. Thr3, at the unstructured amino terminus also exhibits a low *S*^2^ (**Error! Reference source not found**.).

**Table 1:** Methyl Order Parameters for A26I-apo-TRAP, obtained from fitting intensity ratios of peak intensities in single- and double-quantum cross-correlated relaxation experiments [42].

| Resi | Residue | $S^2$ |
| --- | --- | --- |
| 3 | Thr | $0.36 \pm 0.06$ |
| 10 | Ile | $0.56 \pm 0.05$ |
| 20 | Ile | $0.43 \pm 0.02$ |
| 23 | Thr | $0.83 \pm 0.03$ |
| 26 | Ile | $0.24 \pm 0.01$ |
| 28 | Thr | $0.33 \pm 0.01$ |
| 43 | Ile | $0.49 \pm 0.07$ |
| 47 | Thr | $0.27 \pm 0.01$ |
| 50 | Thr | $0.53 \pm 0.07$ |
| 53 | Ile | $0.52 \pm 0.06$ |
| 61 | Ile | $0.68 \pm 0.04$ |
| 63 | Thr | $0.99 \pm 0.12$ |
| 68 | Ile | $0.26 \pm 0.01$ |

### Solid-State NMR Backbone and Side-Chain Resonance Assignments

To extend the backbone and side-chain resonance assignments for apo TRAP we conducted solid-state NMR experiments on uniformly ^13^C,^15^N-labeled wild-type TRAP. Prior solution NMR studies of apo-TRAP, at 55 °C and 800 MHz, yielded resonance assignments for only ∼49% of the protein backbone due to absence of many signals in correlation spectra stemming from conformational exchange. While solution NMR experiments were conducted at elevated temperatures to increase the overall molecular tumbling rate and obtain narrower resonance widths[24,25], because linewidths in solid-state NMR experiments are independent of the rotational correlation time[28] we reasoned that by recording NMR spectra in the solid-state at lower temperatures we may be able to freeze out motions that lead to broadening and disappearance of NMR signals [37]. Specifically, we recorded a set of cross-polarization (CP) based triple-resonance experiments on apo TRAP at 4 °C and 800 MHz, including 3D NCACX, NCOCX, and CANCO[44], to establish sequential resonance assignments. Comparison of the ^15^N-^13^C projections from solution-state TROSY-HNCA recorded at 55 °C and solid-state NCACX spectra show good agreement (Figure 6) with average Δδ for ^15^N amide and ^13^Cα shifts of 0.16 and 0.07 ppm, respectively. In addition, DARR spectra recorded at 45 °C show minimal chemical shift perturbations compared to 4 °C, indicating that the structure of the rigid core remains unperturbed, consistent with the above noted comparisons of solid-state and solution state NMR spectra recorded at 4 °C and 55 °C (Figure S 4).

**Figure 6.**
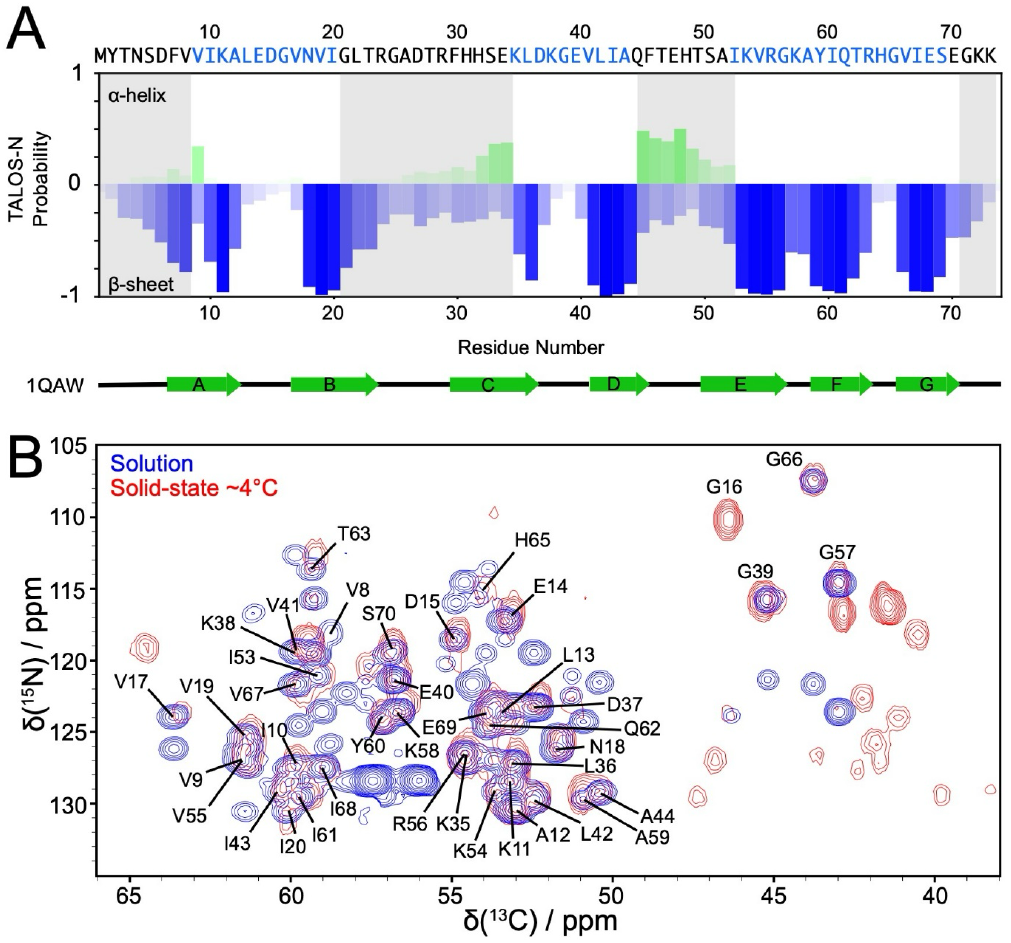
Solid state NMR data of apo TRAP are consistent with solution measurements. (A) TALOS-N secondary structure prediction from chemical shifts assigned in solid-state at 4°C. Primary sequence above with annotation of secondary structure observed in holo TRAP crystal structure. (B) Solution- and solid-state NMR spectra are consistent with local dynamic disorder. Overlay of solution 800 MHz TROSY-HNCA spectrum (blue) with 2D projection from the 3D solid-state MAS NCACX spectrum (red) recorded at the indicated temperatures. Backbone resonance assignments as indicated, unlabeled blue signals correspond to i-1 correlations and unlabeled red signals correspond to sidechain chemical shifts. Close agreement between solution and solid-state spectra indicates similar structures under both conditions. Additional signals in the solid-state spectrum are from aliased Lys/Arg sidechain resonances or C^β^/C^γ^ correlations from sidechain amides of Asn and Gln. Line widths for ^13^C in the NCACX are ∼1 ppm, indicative of a well folded protein. Approximately 50% of possible N-C^α^ correlations are visible in the solid-state experiment.

Backbone ^13^C and ^15^N resonance assignments could be unambiguously established from the NCACX, NCOCX, and CANCO spectra for 40 out of 74 residues (Figure 6). Additionally, these spectra enabled assignment of several side chain ^13^C resonances that were not accessible from solution NMR data. Chemical shift-based secondary structure prediction with TALOS-N[45] shows that the residues observed in CP-based experiments adopt the same secondary structure that is seen in crystals of holo-TRAP. Signals for residues 9-20, 36-44, and 53-70 all were assigned in solid-state spectra of apo TRAP and are predicted from their shifts to adopt either a β-strand or loop conformations.

The CP-based solid-state experiments recorded at 4 °C reveal that dynamic disorder persists at the reduced temperature. Notably, several signals expected for the 74-residue protein are absent in the C^α^/C^β^ region of a 2D ^13^C-^13^C DARR spectrum (Figure 7B). For example, only one threonine (Thr63) out of the six exhibits detectable C^α^-C^β^ and C^β^-C^γ^ correlations, and in the alanine region only three of the five correlations are observed, the other correlation in that region arises from a leucine side chain. Aromatic sidechain correlations are observed only for His65 and Tyr60 in the rigid core on β-strand F (Figure 7B).

**Figure 7.**
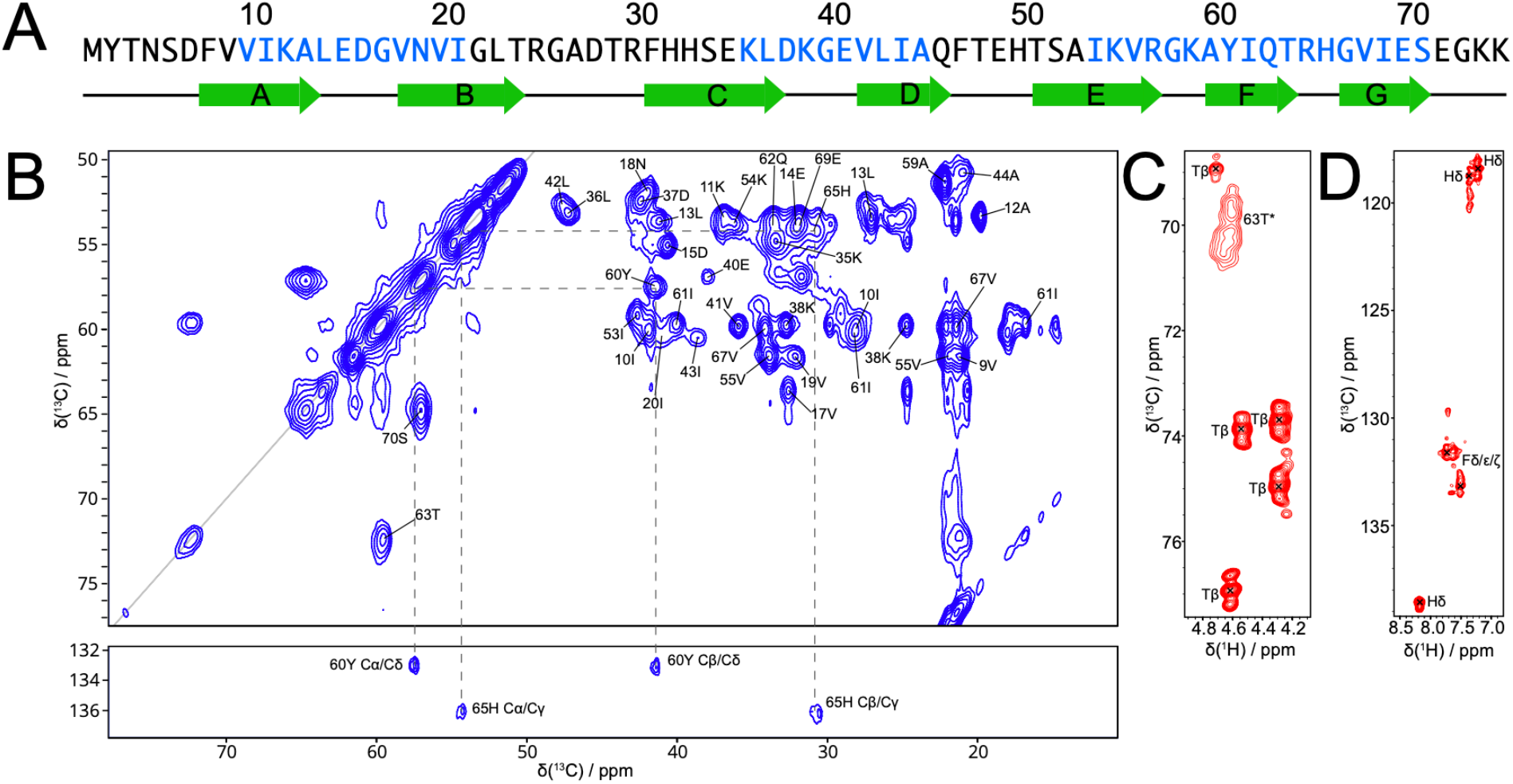
CP- and INEPT-based spectra distinguish structured and flexible regions of apo TRAP. (A) Primary sequence of TRAP. Residues in blue produced signals in solids CP-based experiments. (B) ^13^C-^13^C DARR spectrum with 50 ms mixing time at 11 kHz MAS, 800 MHz field [U-^13^C, ^15^N]-apo-TRAP. The alanine C^α^-C^β^ region only shows three of the five alanine resonances (12, 44, 59). Correlations are observed for only one of the six Thr, Thr63. (C) ^13^C-detected 2D INEPT spectrum shows five distinct sets of Thr Cβ-Hβ correlations, with typical one bond C^α^-C^β^ J-coupling values of ∼34 Hz. The broad peak 63T* corresponds a minor state of the Cβ-Hβ correlation of Thr63 seen in panel B. (D) Aromatic region of the 2D INEPT showing histidine and phenylalanine sidechain correlations.

Given that DARR, NCACX, NCOCX, and CANCO experiments rely on dipolar coupling-based magnetization transfers, the assigned regions correspond to the rigid core of apo TRAP. In addition, several other residues in the structured regions of holo TRAP did not have observable resonances in the CP-based solid-state NMR spectra of apo TRAP. Residues 30-34 in the C-strand, 23-28 in the BC loop, and 46-49 in the DE loop, were “invisible” in conventional CP-based experiments; their absence likely arises from the motional averaging of the dipole-dipole couplings that are required for magnetization transfer[46]. To complement the CP-based solid-state NMR experiments described above and extend our NMR analysis to the flexible regions of apo TRAP we used INEPT-based NMR experiments recorded under MAS which are mediated by scalar (*J*) coupling interactions not averaged by isotropic or near-isotropic local motions[47]. This permits observation of signals arising from the most dynamic regions of the protein while ‘filtering out’ signals arising from the rigid core residues. ^13^C-detected refocused INEPT[48] experiments show an increase in signal from 4°C to 45°C, consistent with an increase in local protein dynamics (Figure S 5).

The INEPT-based ^1^H-^13^C correlation spectrum recorded at 45 °C contains a number of intense correlations from the aromatic and aliphatic regions of residues such as Thr, His, and Phe located outside of the rigid core (Figure 7 C,D). Specifically, this spectrum shows five signals with H^β^ and C^β^ chemical shifts that can only be attributed to Thr residues, with ∼35 Hz ^13^C J-couplings[49]. There is an additional signal at ∼70 ppm that is broadened compared to the other Thr resonances which we assign to Thr63, consistent with broadening that is also seen in the DARR spectrum (Figure 7B). The aromatic region of the spectrum reveals ^1^H-^13^C correlations representative of His and Phe sidechain carbons (120 ppm and 130 ppm, respectively). These aromatic signals can only come from residues 30-32 in the C-strand or 46 and 49 in the DE loop, supporting the conclusion that the C-strand undergoes significant motions in apo TRAP.

### Apo TRAP Sidechain Rotamers

The chemical shifts of C^α^, C^β^, C^γ1^, C^γ2^, and C^δ^ nuclei from all Ile residues (10, 20, 43, 53, 61, and 68) in both apo and holo TRAP were assigned from 2D and 3D solid-state NMR spectra [44] (Figure S 6). Spectra for holo TRAP, recorded at 1.2 GHz, were sufficiently resolved to enable assignment from 2D spectra alone, while 3D spectra were required for assignment of apo TRAP, recorded at 800 MHz. These assignments were used to predict the populations of the {χ_1_, χ_2_} rotameric states by reference to chemical shifts predicted for discrete rotameric states [50] in the fast-exchange regime [51]. Based on these data, in apo TRAP Ile residues 10, 20, 43, 53, and 61 are predicted to sample at least two rotameric states (Figure 8) with the major state (having a probability >0.5) being {*g*_*-*_*/t*} (gauche-/trans). Ile 20, 43, and 53 exchange between three separate states ({*g*_-_/*t*}, {*g*_+_/*t*}, and {*g*_-_/*g*_-_}), whereas Ile 10 and 61 only exchange between two states. Comparing the rotamer distributions predicted for apo TRAP with those predicted for holo TRAP shows modest changes in the rotamer distributions. For Ile43, the most dominant rotameric state shifts from {*g*_-_,*t*} to {*g*_+_,*t*}, while Ile53, with which it pairs in the inter-protomer interface sees a small increase in its favored {*g*_-_,*t*} state. By comparison, in holo TRAP in crystals (1QAW, 1C9S, 1GTF, with resolutions down to 1.75 Å), all Ile residues are modeled as adopting the {*g*_-_,*t*}, except Ile68 which was modeled in the {*t,t*} conformation [19]. The ligand-induced chemical shift changes thus illuminate population shifts arising from binding-coupled changes in the free energy landscape of the protein.

**Figure 8.**
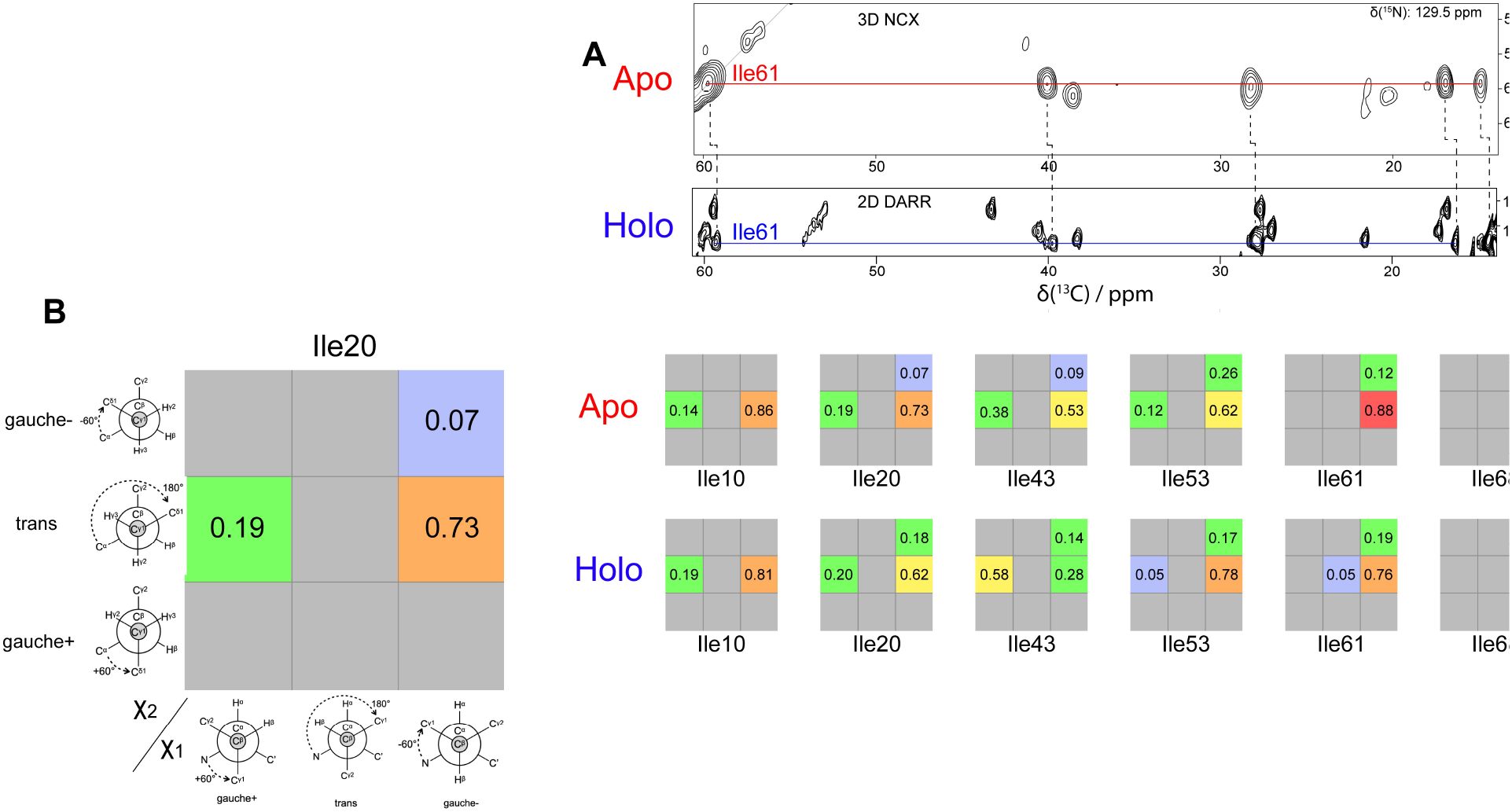
Trp binding results in altered rotamer distributions for Ile side chains. (A) Top, 2D slice from NCACX spectrum collected at 800 MHz of apo TRAP. Red-line passes through cross-peaks that arise from C^α^ correlations to the side chain carbons. Bottom, strip from DARR spectrum collected at 1.2 GHz of holo TRAP. Blue-line passes through cross-peaks that arise from C^δ^ correlations to the rest of the side chain. (B) Left, schematic of rotamer distribution graph. Right, rotamer distributions for Ile residues were calculated from the sidechain ^13^C chemical shifts.

## Discussion

This study builds upon prior solution NMR studies of TRAP [24,25], a 91 kDa homo-oligomeric ligand-activated gene-regulatory protein, with the goal of advancing a mechanistic understanding of its activation by Trp. Introduction of Thr methyl labeling allowed us to probe the dynamics of residues directly responsible for chelating the Trp ligand. Application of solid-state NMR methods allowed us to extend resonance assignments of apo TRAP thereby illuminating how ligand-coupled conformational changes influence subsequent binding events, giving rise to its positive homotropic cooperativity[21,22].

### Concerted motions gate Trp binding

Internal motions in apo TRAP allow it to transiently sample conformations that may facilitate Trp binding. Most of the methyl groups in apo TRAP exhibit relaxation dispersion in ^13^C CPMG experiments (Figure 4, **Error! Reference source not found**.) [25], indicating the presence of dynamics on the µs-ms time scale. In prior NMR relaxation experiments conducted with ILV-methyl labeled A26I-TRAP [25], the single probe on the BC loop, Ile26, exhibited fast exchange dynamics, but the remaining methyl RD curves from Ile, Leu and Val residues in the core of the protein could not be reasonably fit with a global two-state exchange process. Here we find that RD curves for all six Thr γ_2_ methyls and three of the Ile δ_1_ methyls could be fit with a global two-state exchange model with an exchange rate *k*_ex_ of 1700 s^-1^. With Δ*ω* for ^13^C methyl groups not exceeding ∼200 Hz (∼1 ppm at 800 MHz), the fitted exchange rate is in the fast limit, convoluting the minor state populations and their chemical shifts. Nevertheless, since the Trp binding sites are occluded in the Trp-bound crystal structures and the rate of Trp binding to TRAP is vastly slower than the diffusion limit [19,52], we interpret exchange process to reflect concerted sampling of a transiently populated “open” conformation of the binding loops, which could permit entry of Trp (Figure 9).

**Figure 9.**
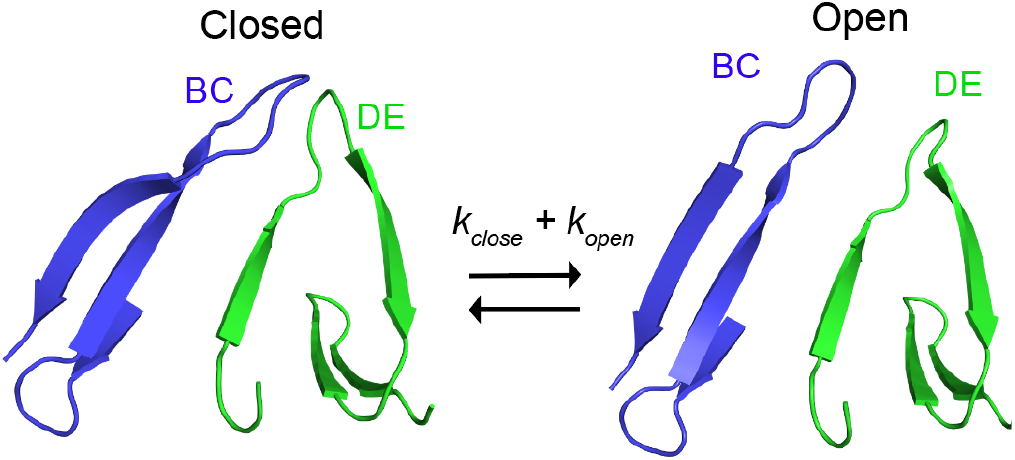
Concerted conformational exchange provides a mechanism for gating access to Trp binding sites. In the apo-state BC/DE loops exchange between an open and closed (holo-like) conformation independently from one another. Successful binding can only occur when both loops are in the open state as the closed (holo-like) state completely shields the cavity. After the first binding event occurs the adjacent protomer’s BC/DE loop favors the open conformation allowing for subsequent binding events to occur more rapidly.

### The Free Energy Landscape of Apo TRAP is Rough and Broad

Solid-state NMR and solution NMR relaxation measurements expand our understanding of dynamics in apo TRAP. NMR relaxation and chemical shift data indicate that Ile and Thr sidechains in apo TRAP sample multiple conformations over both the intermediate µs-ms and fast ps-ns time scales. As noted above, methyl relaxation dispersion data provided evidence for concerted fluctuations in the Trp binding loops that gate access by the ligand. The other Ile methyl probes in apo TRAP also exhibit RD profiles, indicative of extensive conformational sampling throughout the protein.

Cross-correlated methyl relaxation NMR experiments carried out (Figure 5, **Error! Reference source not found**.) show that apo TRAP is also dynamic on the ps-ns timescale. Apo A26I-TRAP has many methyl groups with low *S*^2^ order parameters indicative of fast methyl axis reorientation. Moreover, Ile and Thr methyl groups exhibit dramatically different *S*^2^ values, despite exhibiting similar µs-ms motions (Figure S 7). Thr63 is particularly notable as it could be globally fit to the same two-state exchange process as the other Thr residues but exhibits a high order parameter (S^2^ = 0.99) indicative of rigidity on the timescale of molecular tumbling. This is also consistent with the unique appearance of the Thr63 methyl signal in both the CP-based and INEPT-based solid state NMR spectra (Figure 7). This residue is very highly conserved across bacterial species[53], and in crystal structures the sidechain Oγ_2_ is observed to hydrogen bond to the amide of His65 in the tight βFG turn[19,54], so its rigidity on the fast timescales likely reflects its role in stabilizing local fold of the protein.

The solid-state NMR spectra provide further support for a conformationally variant apo TRAP protein. Cross-polarization NMR methods make use of strong dipolar couplings to transfer magnetization between spins and generate correlations in multidimensional spectra [55,56]. Internal motions result in reorientation of dipoles relative to the static magnetic field, which in turn leads to averaging of the dipolar couplings and loss of signal in CP-based spectra. Conversely, the same rapid dynamics enables preservation of transverse magnetization and observation of signals in INEPT-based spectra of solids[47]. As such, the observation of intense signals in MAS INEPT-based spectra corresponding to the Thr residues in the BC and DE loops, as well as for the His and Phe residues in the DE loop (Figure 7), is strongly suggestive of those regions of the protein being flexible, even at 4 °C in the solid state. In total, backbone resonance assignments could be obtained by solid-state NMR for 40 out of the total 74 residues of apo TRAP; these are consistent with those obtained previously from solution experiments at 55 °C and correspond to the well-structured rigid core of the protein rings. Absence from the CP-based spectra of signals from the amino terminus, BC and DE loops, and C-terminal residues (Figure 6), is indicative of their disorder and flexibility in the solid state.

### Trp Binding Results in a Shift in the Free Energy Landscape

Chemical shift data provide rich structural information about differences between apo and holo TRAP. Comparison of methyl spectra of A26I-TRAP in the absence and presence of Trp (Figure 2) reveal perturbations in both the ^1^H and ^13^C shifts, indicative of changes in the local environment experienced by these methyl probes. In the absence of Trp four of the six Thr methyl resonances have nearly degenerate ^13^C chemical shifts (∼21.5 ppm), while Trp binding disperses these resonances ∼1 ppm (Figure 2). We interpret these perturbations as arising primarily from changes in the time-averaged rotameric states adopted by those residues[57].

NMR resonance assignments of Ile sidechains allowed interrogation of rotamer populations in apo an holo TRAP (Figure 8). Population distributions predicted from chemical shifts generally agree with those modeled in the holo TRAP crystal structures. In those structures, from crystals that diffracted down to a resolution of 1.75 Å, all Ile sidechain χ_1_ and χ_2_ torsion angles were observed to adopt {*g*_*-*_,*t*} configurations, except for Ile68 which is present as the {*t,t*} rotamer[19,54]. Comparing the ^13^C chemical shifts of the Ile sidechains from solid-state NMR spectra of apo and holo TRAP (Figure 8) suggests that Trp binding only modestly alters the rotamer distributions. Nevertheless, even modest shifts in the free energy landscape can contribute significantly to the thermodynamics of ligand response [58].

### Cooperativity from Population Shifts Upon Initial Trp Binding

Positive thermodynamic cooperativity dictates that after one ligand binds, subsequent binding events may occur with higher affinity, as measured by the equilibrium constant *K* between the free and ligand-bound states. This cooperativity can be quantified from the free energy difference between binding events ΔΔ*G* = -*RT* ln *α*, where *α* = *K*’/*K* is the observed fold change in affinity for subsequent compared to initial binding. Ligand titrations monitored by calorimetry and native mass spectrometry, and analyzed with a nearest-neighbor Ising model [21,22], reveal that Trp binding to TRAP is modestly cooperative with ΔΔG coupling energies between adjacent sites of about -1 kcal mol^-1^, or ca. -1.7 *RT*. Understanding the basis for Trp-Trp cooperativity in TRAP requires identifying the origins of this thermodynamic difference between initial and subsequent binding events.

We propose that Trp-Trp cooperativity in TRAP results from reduced conformational sampling (dynamics) at adjacent sites upon initial biding, driving a population shift that favors subsequent ligand binding events (Figure 10). Considering the free energy landscape of an individual site, this can be described as rough and shallow, with many different conformations being similarly populated, including one that resembles the bound state. Binding of a ligand to a site induces rigidification of the BC and DE loops, which also result in reduced conformational sampling of the adjacent sites. In addition, ligand binding results in modest shifts in the rotamer populations of the side chains in the core of the protein that may also propagate from one site to the next.

**Figure 10.**
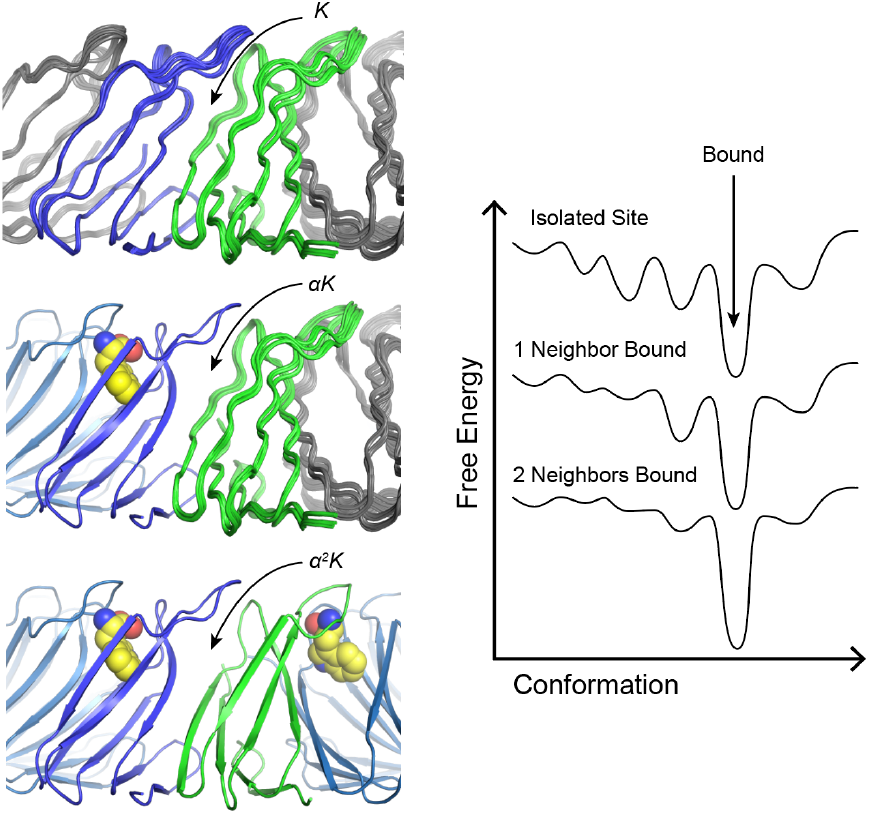
Model for conformational shift in Trp-Trp cooperativity in TRAP. For an isolated Trp binding site with no bound neighbors, the free energy landscape is shallow and rough, with many conformational states being populated. The binding equilibrium constant K reflects the free energy penalty of generating order in the site. Binding of an adjacent Trp reduces conformational sampling on one side of the binding site and narrows the range of accessible conformations, resulting in a more favorable affinity αK. Having two adjacent Trp site occupied further narrows the free energy landscape, resulting in more favorable Trp binding.

This model is supported by several observations from the current study. First, as measured by ^13^C RD, ^1^H-^1^H cross-correlated relaxation experiments, and solid-state CP and INEPT-based NMR experiments, the Trp binding sites in apo TRAP feature a high degree of temporal disorder (dynamics). RD profiles from Ile and Thr methyl groups exhibit exchange on ms time scales, and much of these dynamics are quenched upon Trp binding. Order parameters for the Ile and Thr residues for apo TRAP are low compared to other characterized globular proteins, whereas reduction in those degrees of freedom upon ligand binding would necessarily incur an entropic penalty that would have to be balanced by other favorable interaction[8,59–62]. Residues Thr 23, 28, and 47 have particularly low S^2^ values (<0.4) in apo TRAP, which is indicative of a high degree of conformational entropy[61,63]. Thus, allosteric activation of TRAP by Trp could involve extensive conformational restriction, which is further supported by the large negative heat capacity change ΔC_p_ induced by Trp binding[22]. The contribution of conformational entropy changes to total binding entropy change can in principle be estimated from changes in NMR order parameters[15,62]. Here we assume that Trp binding produces an average methyl-order parameters of 0.8 for Thr and 0.6 for Ile methyls, that methyl axis motion can be approximated by diffusion in a cone, and that increase in the *S*^2^ arises from reduction in concerted motion to estimate the change in protein conformational entropy (Δ*S*_conf_) associated with binding[64]. These assumptions result in an entropic cost for ordering a site of ca. +7 *RT* (+4.3 kcal mol^-1^ at 40 °C) that would have to be compensated for by favorable changes in enthalpy and/or solvent entropy. This value is on the same scale, though opposite sign, as the calorimetrically-measured ΔS_coupling_ of +8 *RT* (+5 kcal/mol)[22]. Additional entropic costs from conformational changes would be expected from redistribution of the Ile rotamer populations, although most populations changed by <0.2 (Figure 8).

### Conclusion

Overall, these findings support a model where protein conformational dynamics play an important role in Trp binding and Trp-Trp allostery in TRAP. The methyl relaxation dispersion data are consistent with apo TRAP experiencing correlated motions, sampling conformations that could enable Trp binding. In contrast, holo TRAP exhibits quenched dynamics and is characterized by restricted internal motions on the µs-ms time scale. Given that ligand binding is accompanied by local protein restriction, this implies an unfavorable conformational entropy change. If initial binding results in reduction in conformational sampling of neighboring sites, subsequent binding events are accompanied by a smaller ΔS penalty. This conformational narrowing (reduced dynamics) provides a mechanistic explanation for the observed NN cooperativity in Trp binding in TRAP.

## Methods

### Sample Preparation

NMR experiments were performed with either wild-type (WT) *Gst* TRAP, and a point mutant in which Ala26 was replaced with Ile (A26I-TRAP) to provide an additional methyl probe on the BC loop. For each sample, data were recorded in the absence and presence of Trp. NMR samples were recombinantly expressed from *Escherichia coli* grown in defined minimal media. Trp-free apo TRAP was purified under denaturing conditions to ensure complete removal of Trp, as described previously. Solid-state NMR spectra were recorded with U-[^1^H, ^13^C, ^15^N]-TRAP (WT), while the Thr and Ile [^13^CH_3_], U-[^2^H,^15^N]-A26I-TRAP was used for solution-state spin relaxation measurements and HCH/CCH methyl-methyl NOESY for assignments. Isotopic labeling of the sample used for solid-state NMR studies was accomplished by protein expression in *E. coli* BL21(DE3) cells grown on M9 minimal media containing ^15^NH_4_Cl and ^13^C-glucose as the sole nitrogen and carbon sources, respectively. Ile and Thr methyl-labeling was accomplished using perdeuterated metabolic precursors of Thr and Ile. A26I-TRAP was used for solution NMR experiments for consistency with prior spin relaxation experiments; samples were 1 mM A26I-TRAP (monomer) in NMR buffer (100 mM NaCl, 50mM NaPO_4_, pH 8.0, 10% D_2_O). WT TRAP was used for solid-state experiments in NMR buffer without D_2_O. For visualization, PyMOL (http://pymol.org) was used to introduce the Ile26 to Ala mutation using either the 1QAW or 1C9S holo structures as templates.

### Solution NMR Spectroscopy

Spectra were recorded on Bruker Avance III HD 600 and 800 MHz spectrometers equipped with HCN z-axis gradient cryo-probes. Sample temperatures were calibrated using a methanol standard and chemical shifts were referenced externally to DSS.

Thr methyl signals of apo A26I TRAP were assigned using two spectra of A26I-TRAP recorded at 55°C: HCH Methyl-Methyl NOESY to correlate a ^1^H-^13^C pair to a neighboring methyl ^1^H, and CCH Methyl-Methyl NOESY to correlate another ^1^H-^13^C pair to neighboring ^13^C spins[35].

For both apo and holo A26I-TRAP multiple-quantum methyl CPMG experiments were carried out at 20 °C. Dispersion curves for apo TRAP were obtained using interleaved CPMG field strengths of 50, 100, 150, 200, 250, 300, 350, 400, 450, 500, 750, 1000, 1500, and 2000 Hz and 50-4000 Hz for holo TRAP.

Dispersions were calculated using R_2_^Eff^ = -ln(I(ν_CPMG_)/I_0_)/T_CPMG_, where I(ν_CPMG_) is the signal intensity from a spectrum with a refocusing duration of T_CPMG_ at a given frequency and I_0_ is the intensity from the spectrum without refocusing pulses. Peak intensities were measured and converted to transverse relaxation rates (R_2_^Eff^) using NMRPipe[65]. The GUARDD software package was used to fit the relaxation dispersions to the Carver-Richards equation that describes the relaxation contribution from chemical exchange (R_ex_) to transverse relaxation (R_2_) by fitting an exchange rate (*k*_ex_) and population (P_A_) for two-state exchange[66]. Errors for fitted parameters were determined by grid searching a five-dimensional parameter space to minimize χ^2^. Dynamics in the fast exchange regime limited the determination of P_A_, |Δ*ω*_C_|, and |Δ*ω*_H_| as any combination of the three can provide equally good fit to *k*_ex_ when assessed by χ^2^ analysis. Methyl probes that fit for similar parameters were fit to a global model.

Methyl order parameters (*S*^2^) for Ile and Thr residues in A26I-TRAP were obtained from measurement of the intensity ratios of double-quantum and single-quantum coherences generated from ^1^H-^1^H dipolar cross-correlated relaxation experiments[42] using a pseudo-3D spectrum. Data were recorded at 55°C at which previous studies established a correlation time *τ*_c_ = 34.2 ns[67], using the following relaxation delays: 0.25, 0.5, 1, 2, 3, 4, 6, 8, 10, 12, 14, 16, 20, 30, 40, 50 ms. Peak intensities were extracted using NMRPipe and residue-specific *S*^2^ values were obtained by fitting intensity ratios in Python using the lmfit library (https://lmfit.github.io/lmfit-py/). Uncertainties were obtained by from Mone-Carlo simulations using the spectral noise as the uncertainty in peak heights.

### Solid-State NMR Spectroscopy

MAS NMR spectra of U-[^13^C,^15^N]-apo-TRAP were recorded on a Bruker Avance III HD 800 MHz spectrometer outfitted with an Efree HCN 3.2 mm MAS probe. Purified TRAP was sedimented in an ultracentrifuge at at 4°C and 100,000 rpm using a TLA 100.3 rotor. The protein precipitate was loaded into a 3.2 mm rotor using home-built tools in a tabletop centrifuge. The MAS frequencies were controlled within ±1 Hz by a Bruker MAS controller. The sample temperature was regulated by a Bruker VT controller and maintained at 4 °C throughout experiments as measured by a KBr external reference.

Typical ^1^H, ^13^C, and ^15^N 90° pulse lengths were 2.5, 5.0, and 6.25 µs, respectively, and ∼90kHz SPINAL-64 proton decoupling was applied during chemical shift evolution periods[68]. 2D ^13^C-^13^C DARR spectra were recorded at 11.111 kHz MAS rate and the following mixing times: 5, 15, 50, 100, and 500 ms, with a 11.111 kHz ^1^H field applied during the mixing time[69]. 2D NCA, 2D NCO, 3D NCACX, 3D NCOCX, 3D CANCO, spectra were recorded at an MAS frequency of 11.111 kHz[69,70]. Band-selective CP (SPECIFIC-CP)[56] was used for magnetization transfer from ^15^N-^13^C in the NCACX and NCOCX, as well as for ^13^C-^15^N magnetization transfers in CANCO with 6ms and 6ms contact times respectively and 15 ms DARR was used for ^13^C-^13^C magnetization transfer in the NCACX and NCOCX experiments. A DARR spectrum of U-[^13^C,^15^N]-Holo-TRAP was recorded on a Bruker Ascend 1.2 GHz spectrometer outfitted with a 3.2 mm HX MAS probe, with sample temperature maintained at 4 °C.

A 2D ^13^C-refocused INEPT spectrum was recorded following the pulse sequence described by Helmus et al. [48,71]. Typical ^1^H and ^13^C 90° pulse lengths were 2.5 µs and 5.0 µs, respectively. Spectra were recorded at 13 kHz MAS and sample temperature of 45 °C, determined from an inversion-recovery experiment performed on a KBr standard.

Resonance assignments for *Gst* TRAP have been deposited to the BMRB, accession XXXX.

## Abbreviations

TRAP: *trp* RNA-binding attenuation protein
CSP: chemical shift perturbation
CP-MAS: cross-polarization magic-angle spinning
TROSY: transverse relaxation optimized spectroscopy
MQ: multiple-quantum
RD: relaxation dispersion
DQC: double-quantum coherence
SQC: single-quantum coherence
DARR: dipolar-assisted rotational resonance

## Acknowledgments

The authors acknowledge the OSU CCIC NMR facility for access to instrumentation and expert support by Chunhua Yuan and Alex Hansen, and Vidhya Sridharan and Tanya Whitmer for advice in solid-state NMR data collection and processing. Paul Gollnick (University at Buffalo) provided reagents and indispensable advice. This work was supported by grants from the NIH (R01-GM120923 to MPF and PG) and from the National Science Foundation (MCB-2303862 to CPJ). Purchase of the 1.2 GHz NMR spectrometer used for some of the later solid-state NMR data collection was made possible from a Mid-scale RI-1 grant from the NSF (DBI-1935913). RM acknowledges support from NIH T32-GM086252. This study made use of NMRbox: National Center for Biomolecular NMR Data Processing and Analysis, a Biomedical Technology Research Resource (BTRR), which is supported by NIH grant P41GM111135 (NIGMS). Melody Holmquist and Vicki Wysocki (OSU) enabled native mass spectrometry samples to assess sample integrity via the Native MS-guided Structural Biology Center funded by NIH RM1GM149374.

