## Supporting Information for "Insights into Ligand-Mediated Activation of an Oligomeric Ring-Shaped Gene-Regulatory Protein from Solution- and Solid-State NMR"

| From | To | Distance (Å) | HCH Cross-Peak | CCH Cross-Peak | Notes |
|---|---|---|---|---|---|
| Ile61 | Thr63 | 7.4 | Yes | Yes | |
| Ile43 | Thr63 | 4.8 | Yes | Yes | |
| Thr63 | Ile61 | 7.4 | Yes | Yes | |
| Ile53 | Thr50 | 4.5 | Yes | Yes | |
| Thr50 | Ile53 | 4.5 | Yes | Yes | |
| Ile20 | Thr28 | 11.2 | No | Yes | |
| Ile26 | Thr23 | 6.9 | Yes | Yes | *No crystal structure of mutation |
| Thr23 | Ile26 | 6.9 | Yes | Yes | *No crystal structure of mutation |
| Thr28 | Ile20 | 11.2 | Yes | No | |
| Thr50 | Thr47 | 5.7 | No | Yes | |

*Table S 1. Methyl-Methyl NOESY Cross peak assignment table.*

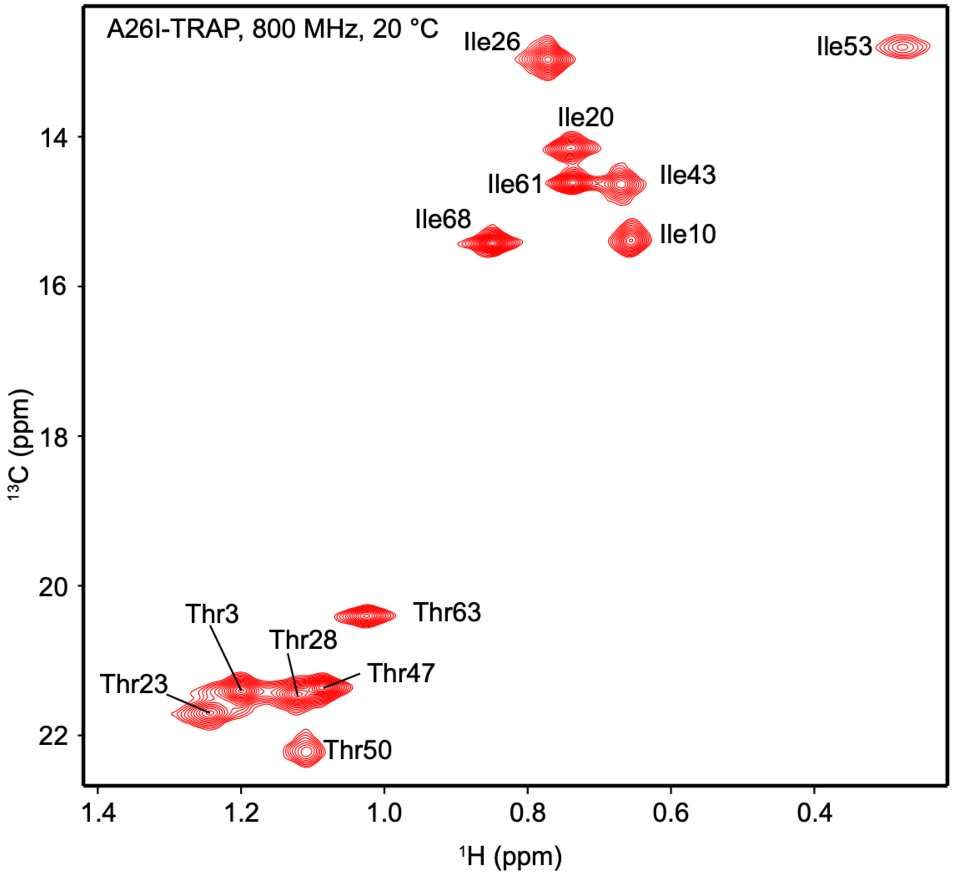


*Figure S 1: TROSY-HMQC of apo [U-$^{2}$H/$^{15}$N, Ile/Thr-methyl $^{13}$C]-A26I-TRAP at 800 MHz, 20 °C. Representative spectrum corresponding to conditions for methyl CPMG relaxation dispersion experiments.*

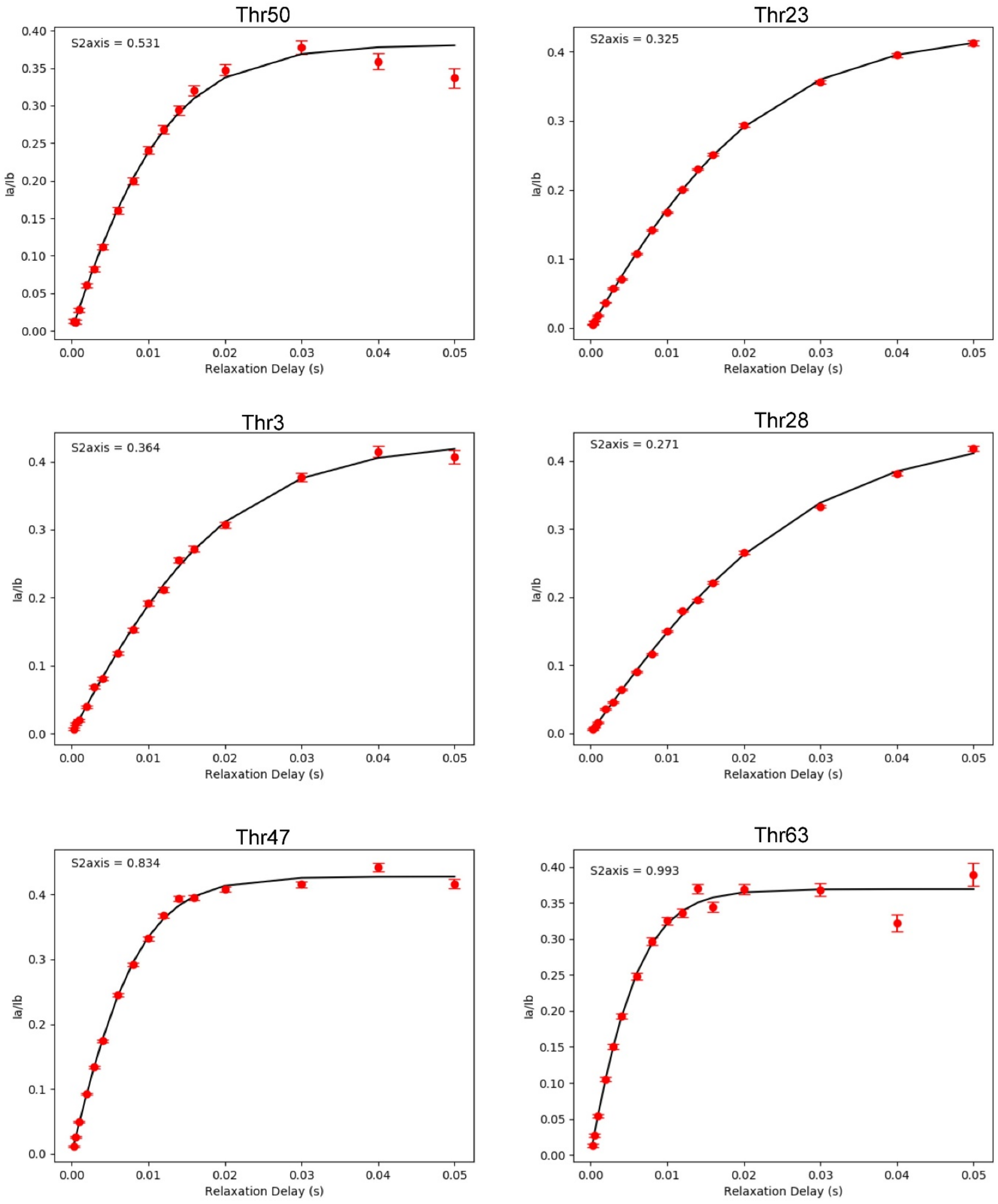


*Figure S 2: $\gamma_2$ Methyl DQC/SQC buildup Curves for Thr residues in apo A26I TRAP.*

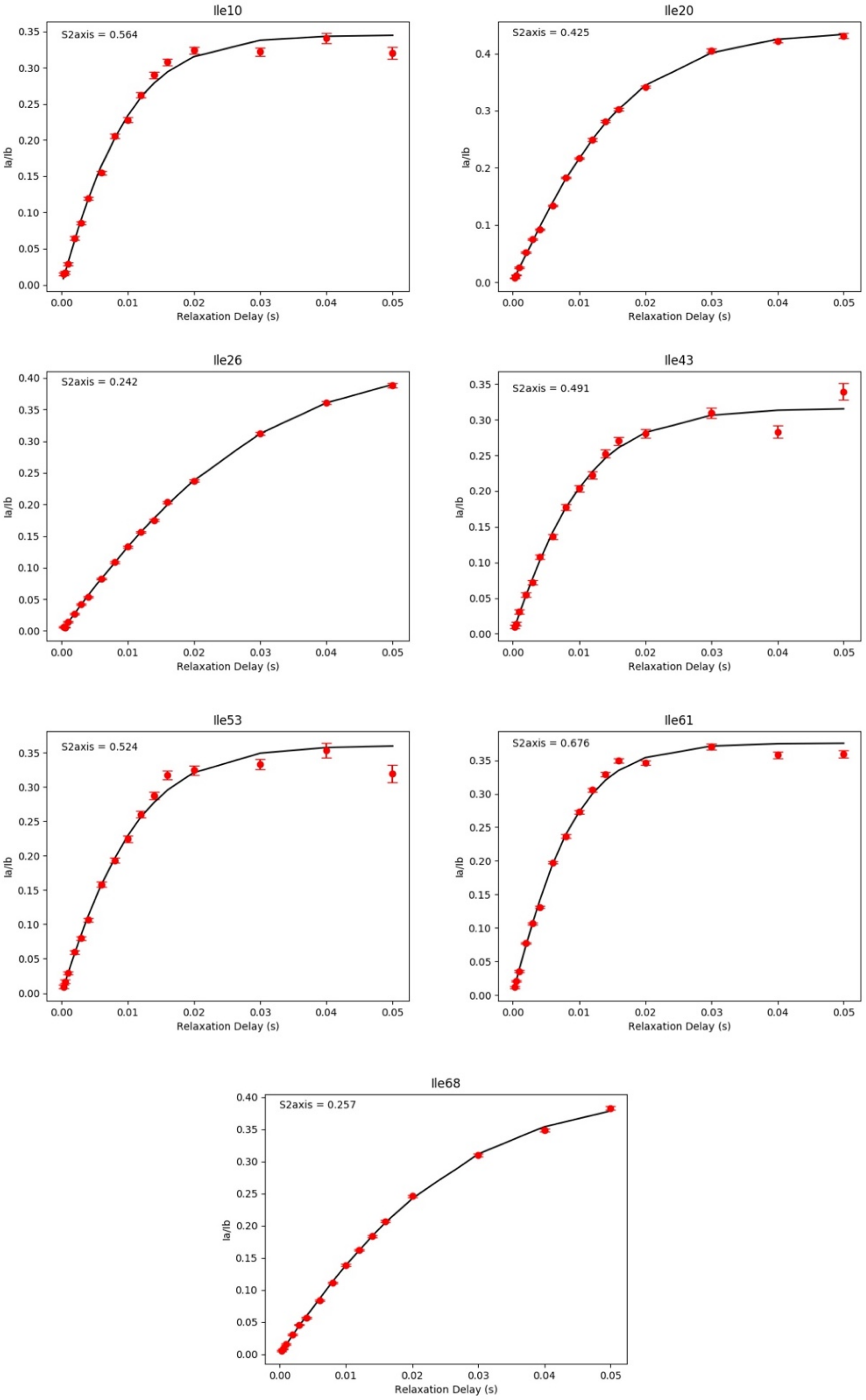


*Figure S 3: $\delta_1$ Methyl DQC/SQC buildup Curves for Ile residues in apo A26I TRAP.*

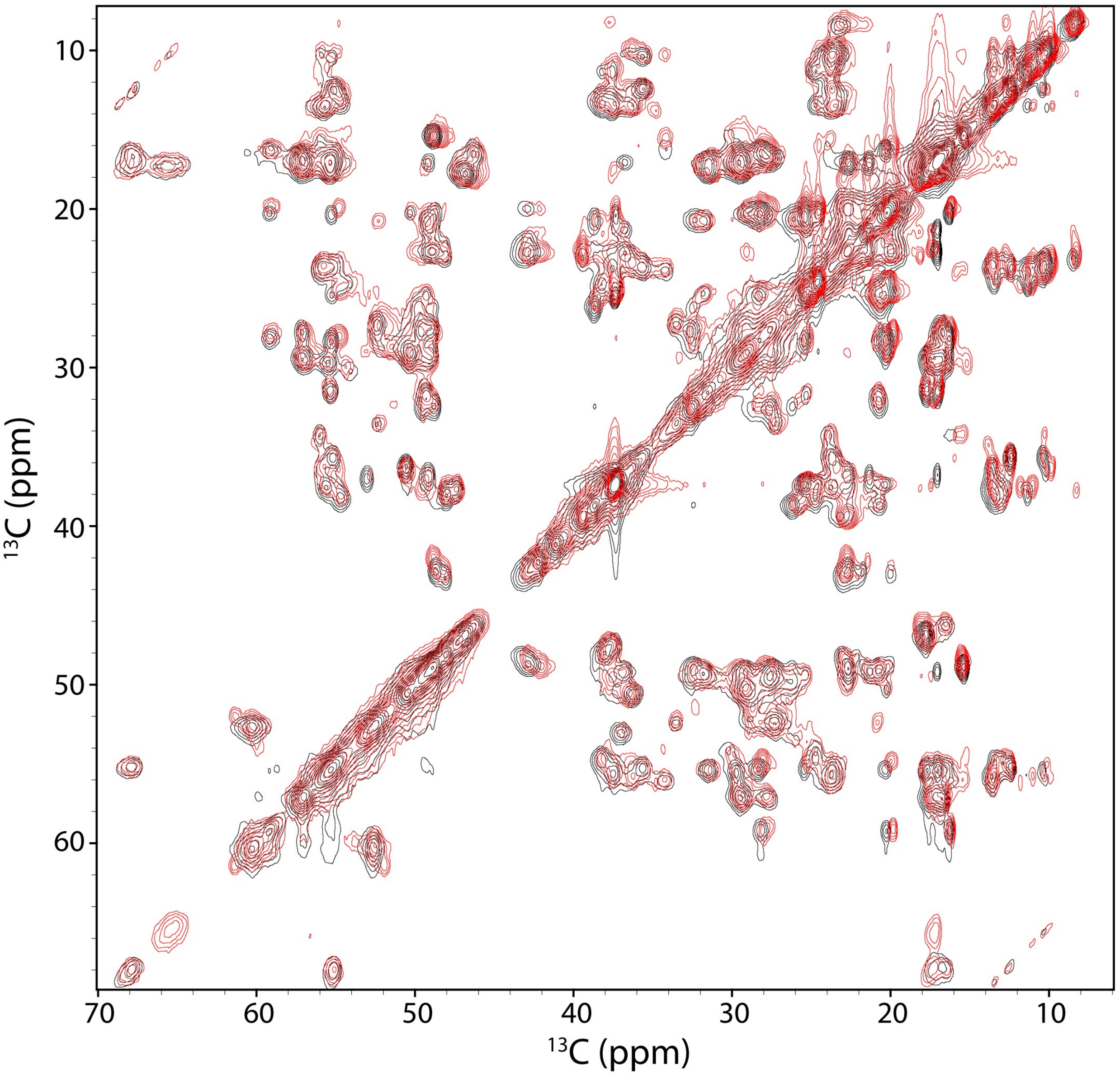


*Figure S 4: Solid-state CP spectra of apo TRAP is not strongly sensitive to temperature. Fingerprint region of the $^{13}$C-$^{13}$C DARR chemical shift correlation spectra of apo TRAP at 4°C (black) and ~45°C (red) using a mixing time of 15 ms at 11.111 kHz MAS speed in a 3.2 mm rotor at 800 MHz.*

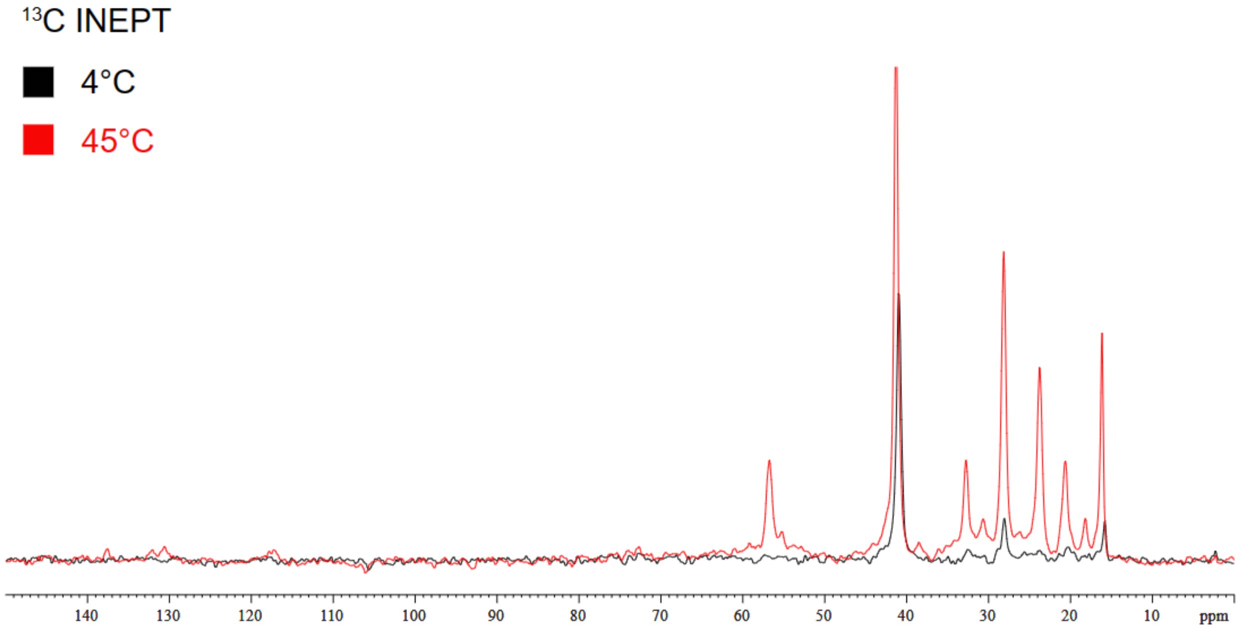


*Figure S 5: 1D $^{13}C$ INEPT spectra of WT apo TRAP at 800 MHz field strength, under a MAS speed of 11.111 kHz in a 3.2 mm rotor at high and low temperatures shows an increase in signal.*

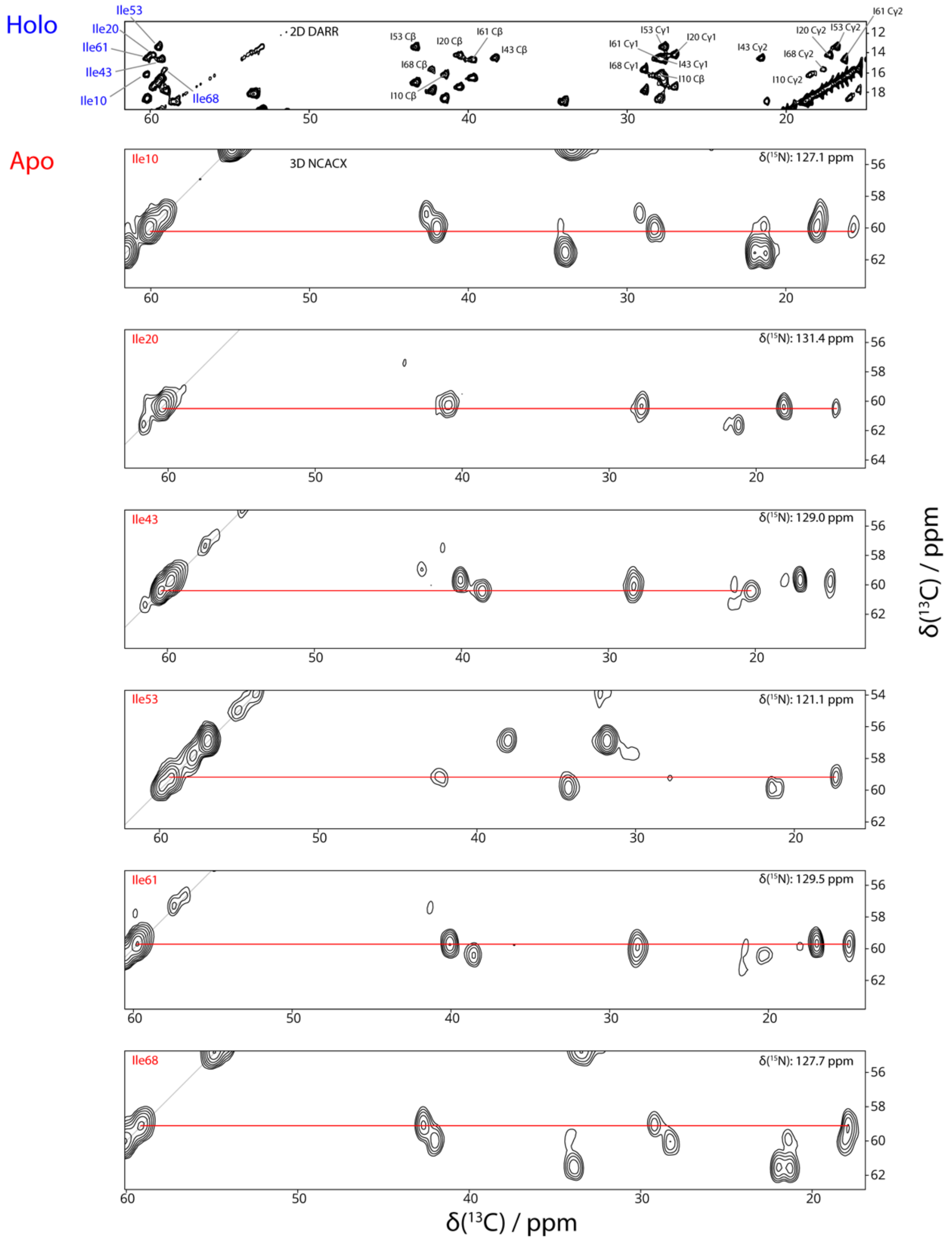


*Figure S 6:Top, $^{13}$C DARR of holo Trp-TRAP recorded with 15 ms mixing time on a 1.2 GHz instrument at 4 °C MAS: 12 kHz. Correlations are observed along the entire side chain, allowing direct $^{13}$C assignments for all Ile in the protein. (For the WT protein residue 26 is Ala, not Ile.) Bottom, strips from the 3D NCACX spectrum of apo TRAP, recorded at 800 MHz, illustrating assignments of the Ile $^{13}$C resonances in apo TRAP. Comparison of the sidechain $^{13}$C shifts in these spectra reveal changes in rotamer distributions.*

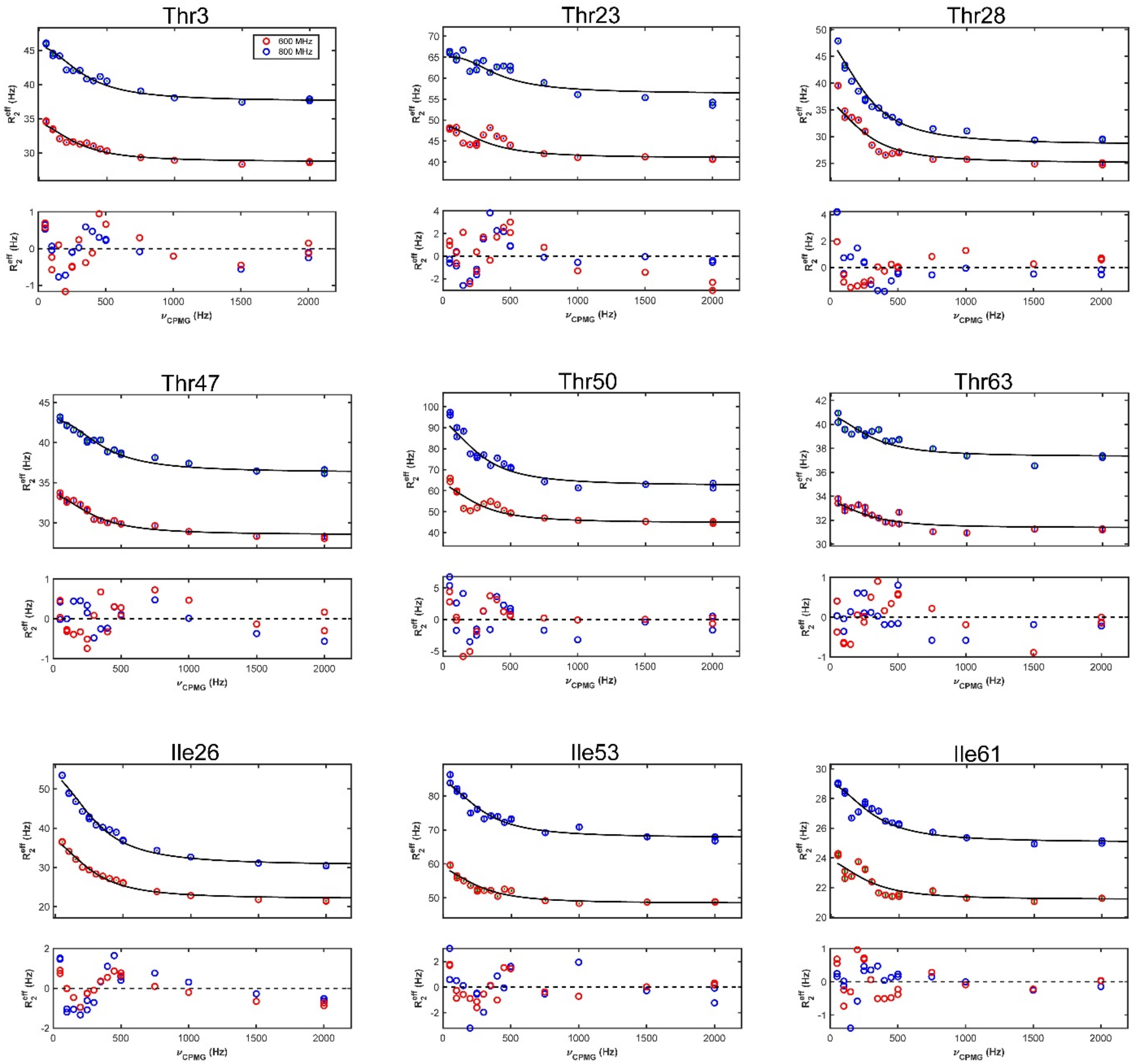


*Figure S 7: Multi-quantum (MQ) methyl RD curves of A26I apo-TRAP at 20°C at 600 (red) and 800 (blue) MHz $B_0$ field strength. The RD curves for all six threonine and three of the Ile δ1 methyl groups (Ile26, Ile53, Ile61) could be globally fit to a single two-state exchange process with a $k_{ex}$ value of 1725 ± 263 $s^{-1}$.*

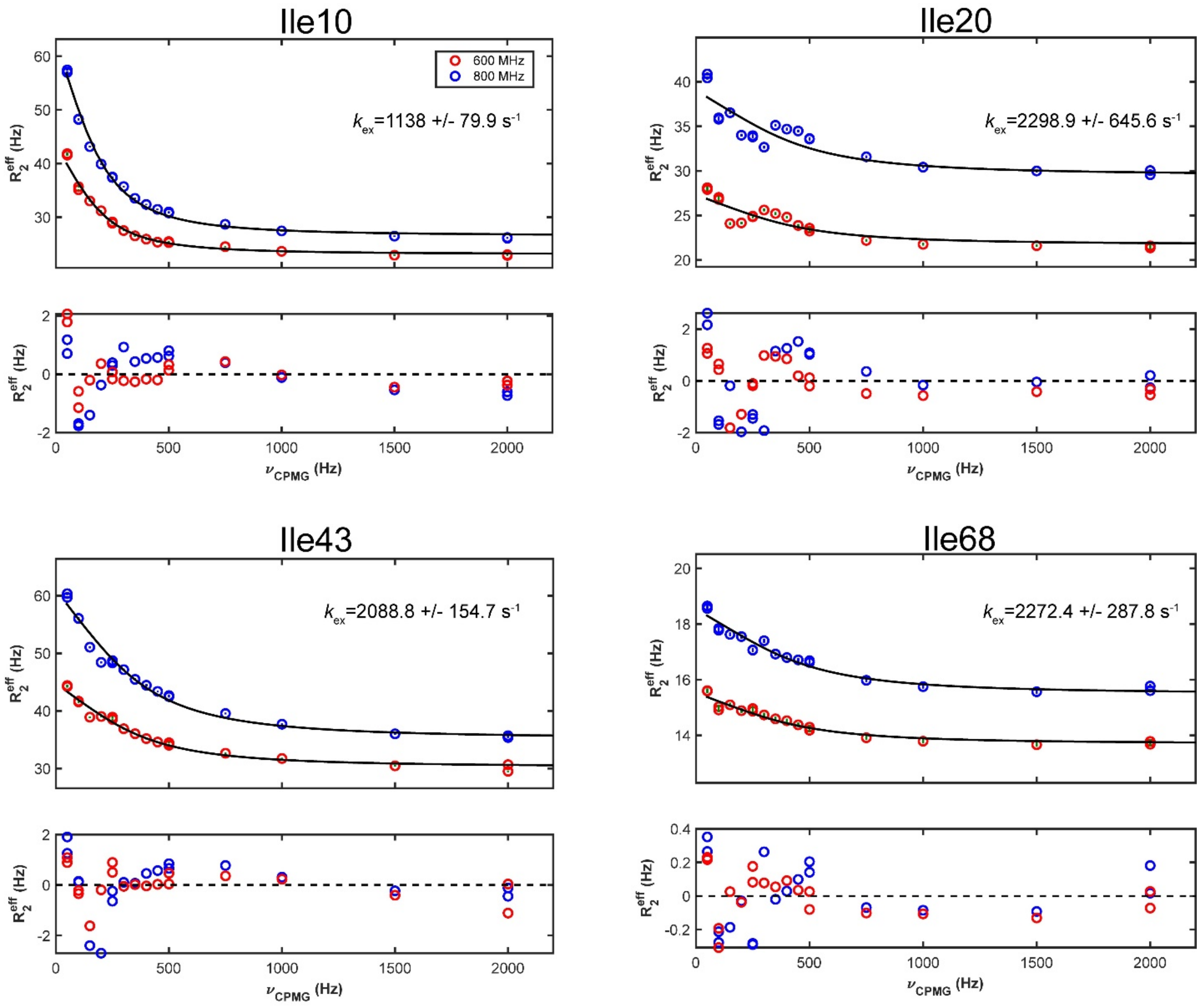


*Figure S 8: Apo A26I-TRAP has several Ile-Cδ$_1$ RD curves that cannot fit to a global model. RD curves here were fit individually with a two-state model and highlight the large difference in μs-ms dynamics with the lowest exchange rate of 1139 $s^{-1}$ and the fastest at 2272 $s^{-1}$.*

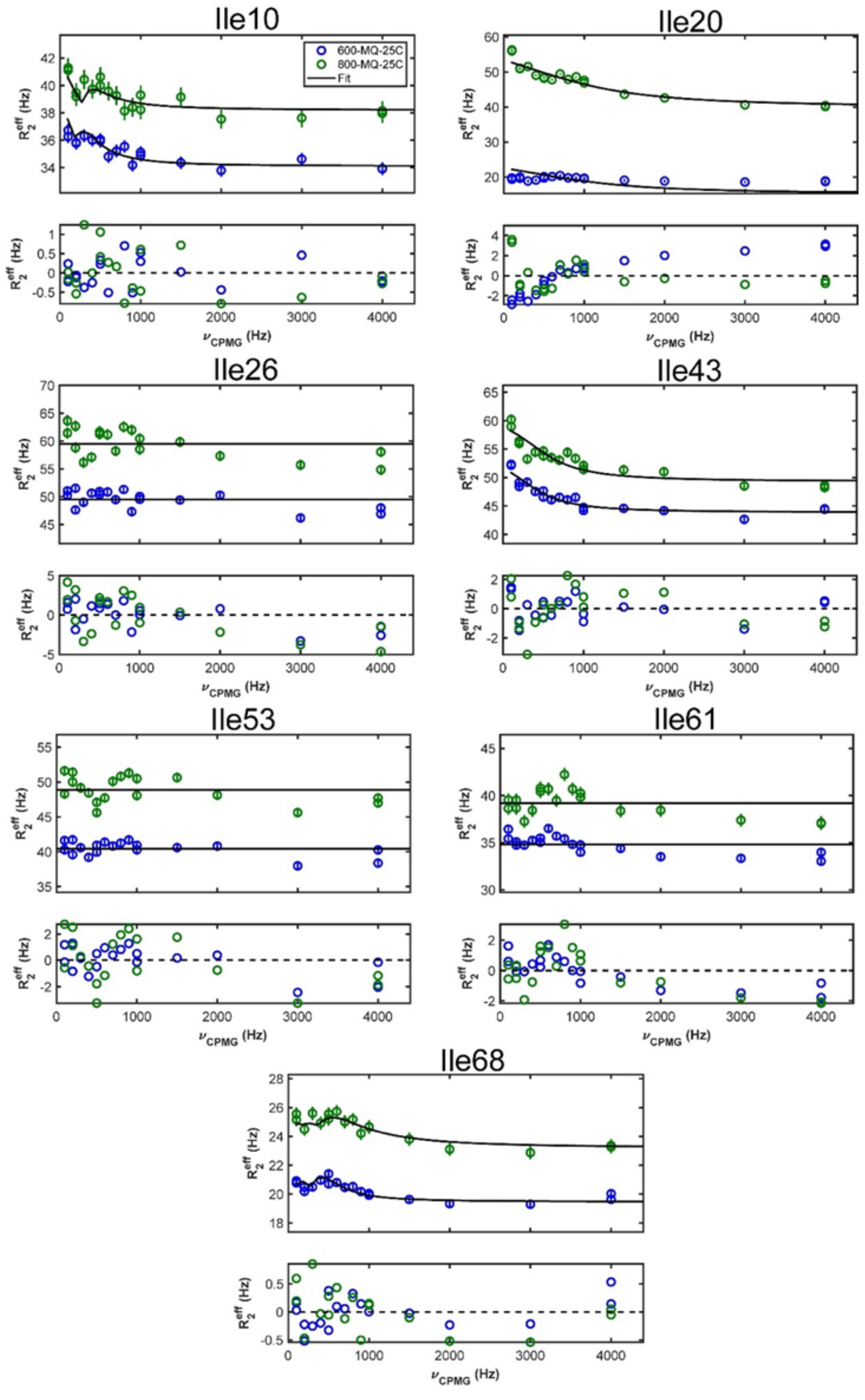


*Figure S 9: Methyl RD curves from A26I holo TRAP collected at 20°C and field strengths of 600 and 800 MHz. Ile residues 26, 53, and 61 show no dispersions, while the remaining residues have significantly reduced exchange rates.*

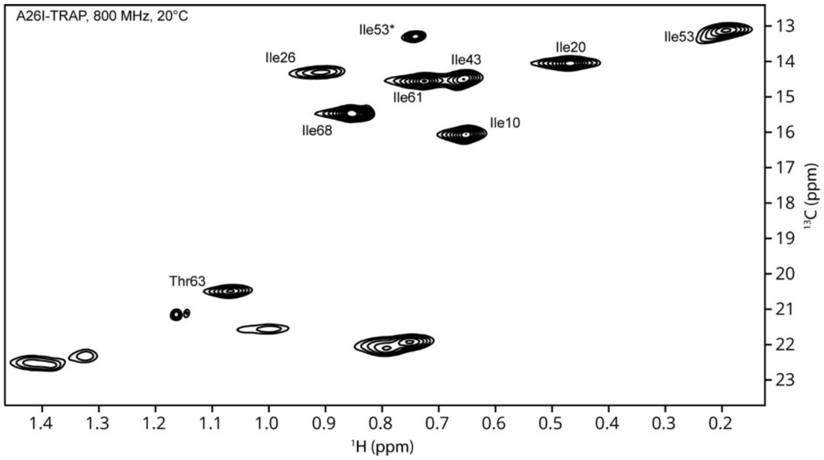


*Figure S 10: Methyl TROSY HMQC of holo A26I-TRAP at 800 MHz and 20°C.*

### Calculation of Entropy from Order Parameters

We calculated the change in conformational entropy from the methyl order parameters using the previously described relationship[64].

$$\frac{\Delta S_p(j)}{k} = \ln\{[3 - (1 + 8 * S_b)^{\frac{1}{2}}]/[3 - (1 + 8 * S_a)^{\frac{1}{2}}]$$

Where $S_b$ and $S_a$ are the order parameters of the methyl bond in states b and a respectively and $\Delta S_p(j)$ is the total rotational entropy change for the methyl bond j. This model assumes that motion occurs as diffusion-in-a-cone. Under the assumption that state changes from a to b occur as concerted motion amongst all Thr/Ile methyl probes, the average $\Delta S_p$ was used to calculate the entropic contributions to free energy ($\Delta S_{conf}$) using the following equation.

$$\Delta S_{conf} = \frac{\Delta S_p(j)}{k} * \frac{N}{k} * 4.18 * T$$

Where $k$ is the Boltzmann constant, N is Avogadro's number, and T is temperature in kelvin, providing the energy in kcal mol$^{-1}$.
